# Atlas of imprinted and allele-specific DNA methylation in the human body

**DOI:** 10.1101/2024.05.01.591988

**Authors:** Jonathan Rosenski, Ayelet Peretz, Judith Magenheim, Netanel Loyfer, Ruth Shemer, Benjamin Glaser, Yuval Dor, Tommy Kaplan

## Abstract

Allele-specific DNA methylation, determined genetically or epigenetically, is involved in gene regulation and underlies multiple pathologies. Yet, our knowledge of this phenomenon is partial, and largely limited to blood lineages. Here, we present a comprehensive atlas of allele-specific DNA methylation, using deep whole-genome sequencing across 39 normal human cell types. We identified 325k genomic regions, covering 6% of the genome and containing 11% of all CpG sites, that show a bimodal distribution of methylated and unmethylated molecules. In 34K of these regions, genetic variations at individual alleles segregate with methylation patterns, thus validating allele-specific methylation. We also identified 460 regions showing parentally-imprinted methylation, the majority of which were not previously reported. Surprisingly, sequence-dependent and parent-dependent methylation patterns are often restricted to specific cell types, revealing unappreciated variation in the human allele-specific methylation across the human body. The atlas provides a resource for studying allele-specific methylation and regulatory mechanisms underlying imprinted expression in specific human cell types.

**Highlights:**

- A comprehensive atlas of allele-specific methylation in primary human cell types
- 325k genomic regions show a bimodal pattern of of hyper- and hypo-methylation of DNA
- Allele-specific methylation at 34k genomic regions
- Tissue-specific effects at known imprinting control regions (ICRs)
- 100s of novel loci exhibiting parentally-imprinted methylation
- Parentally-imprinting methylation is often cell-type-specific

---

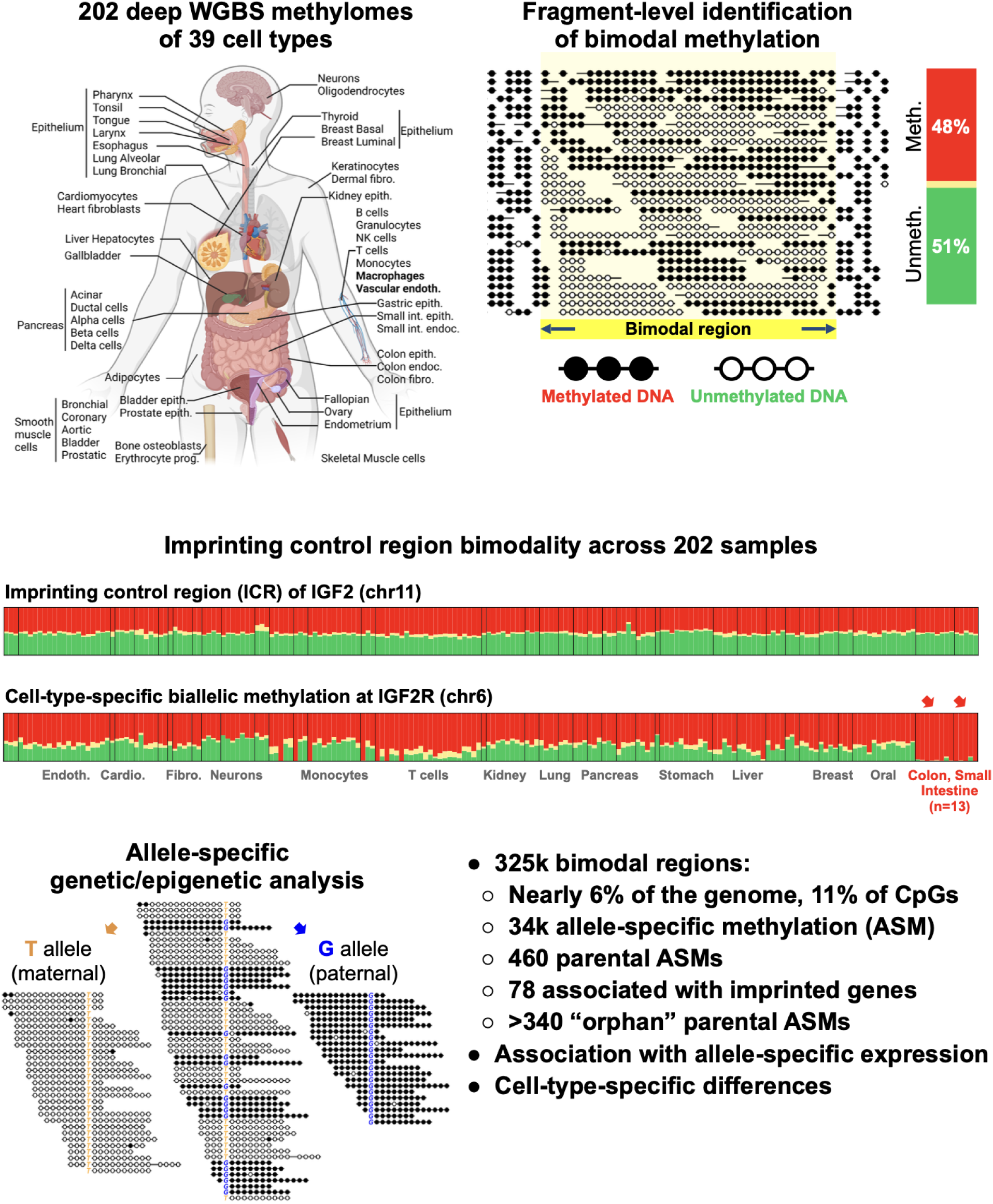

## Introduction

DNA methylation is a stable epigenetic mark that alters the accessibility and 3D packaging of the genome, allowing differentiated cell types to selectively utilize transcriptional programs and maintain their cellular identity throughout life^1^. Methylation patterns are generally identical between the paternal and maternal alleles^2–6^, although a small fraction of the genome, estimated at 5% based on common SNPs, was reported to show allelic methylation differences^4,7–11^.

The molecular basis of monoallelic methylation patterns and their functional consequences vary. In the case of meQTLs, genetic variation is associated with varying levels of methylation. One cis-acting genetic variant is associated with hyper-methylation, whereas another variant is associated with hypo-methylation, possibly to regulate the expression of an adjacent gene^7^. In other cases, for example in mammalian female X-chromosome inactivation, at some early embryonic developmental stage each cell randomly methylates one chromosome, and this selection is then maintained in future cell divisions^8,12–14^. In other cases, allelic methylation relates to the parent of origin, wherein either the maternal or paternal allele is methylated^15–18^. Parental allele-specific methylation differences at imprinting control regions (ICRs), that are established early in the sperm and egg and retained throughout development, serve as the basis for genomic imprinting whereby genes are expressed only from one specific parental allele^19–22^. These epigenetic differences play an important role in placental function and in embryonic development, and dysregulation of imprinted genes is associated with several developmental disorders, including Beckwith-Wiedemann syndrome, Angelman syndrome, and Prader-Willi syndrome^23–28^.

Nonetheless, our understanding of allele-specific methylation remains incomplete. To a large extent, this is due to the fact that most genome-wide methylome datasets are based on DNA methylation arrays (Illumina BeadChip 450K and EPIC), and are limited to a predefined set of CpGs, capturing only 1.5%-3% of the 28M methylation sites in the human genome. Additionally, methylation arrays capture the average methylation levels of individual CpGs, and genetic (SNP) information or epigenetic dependencies between neighboring sites on the same DNA molecule are unobservable. Finally, the study of imprinting and allele-specific methylation in humans was previously limited to few cell types, focusing on easily accessible blood DNA^15, 28–31^.

Several next-generation sequencing studies recently analyzed allele-specific methylation (ASM) from blood^15,29^. Other studies analyzed uniparental disomy samples^30,31^ using a combination of iPSC, ESC and blood cells^32^, or focused on sequence-dependent allele-specific changes^7^. Additionally, allele-specific gene expression was studied across human tissues^16^, using RNA-seq data from GTEx^33^, and reported biallelic expression for known imprinted genes in few tissues (e.g. IGF2 in the liver^16^); similarly tissue-specific allelic expression was reported in mice^34^. Such cases of cell-type-specific escape from parental repression raise questions as to the molecular mechanisms underlying imprinting, their relation to allele-specific methylation, and how they are modified in specific cell types, thus exemplifying the need for a detailed genome-wide pan-tissue atlas of allele-specific methylation. Recently, we characterized the DNA methylation landscape of over 200 surgical and blood samples that were obtained from 135 donors, purified to homogeneity, and deeply sequenced at a whole-genome scale^3^.

Here, we developed computational algorithms for the identification and characterization of allele-specific methylation (ASM), including sequence-dependent effects (e.g. meQTLs) as well as parentally methylated regions. Overall, we identified 325k regions with bimodal DNA methylation patterns, 34k of which overlap heterozygous SNPs that segregate with DNA methylation, thus supporting allele-specific differences. We also identified 460 putative parentally imprinted regions where allele-specific methylation across multiple donors cannot be explained by heterozygous variations. These include 45 known imprinting control regions (out of a total of 55 known ICRs), another 34 novel regions near known imprinted genes and 381 tissue-specific parentally methylated regions, with one novel region validated across 33 parent-child trios. These parental DNA methylation regions are enriched for regulatory regions, polycomb domains, and origins of asynchronous replication. The atlas presented here expands our knowledge of parental imprinting across specific cell types, with implications for understanding the crosstalk between genetic variation, DNA methylation and allele-specific expression.

## Results

To analyze the landscape of allele-specific methylation and imprinting, we revisited our human DNA methylation atlas describing the methylomes of 202 healthy samples, representing ∼40 primary cell types collected and purified from 135 donors^3^. The purity of these samples and the high sequencing depth facilitate a fragment-level analysis of DNA methylation, capturing both genetic and epigenetic information from each sequenced fragment. This dataset offers a unique resource to uncover cell-type-specificity of allele-specific methylation. To further understand how ASM may impact gene expression, we integrated the methylation atlas with allele-specific expression data from GTEx^35^ spanning >50 tissues and cell types (Figure 1).

**Figure 1.**
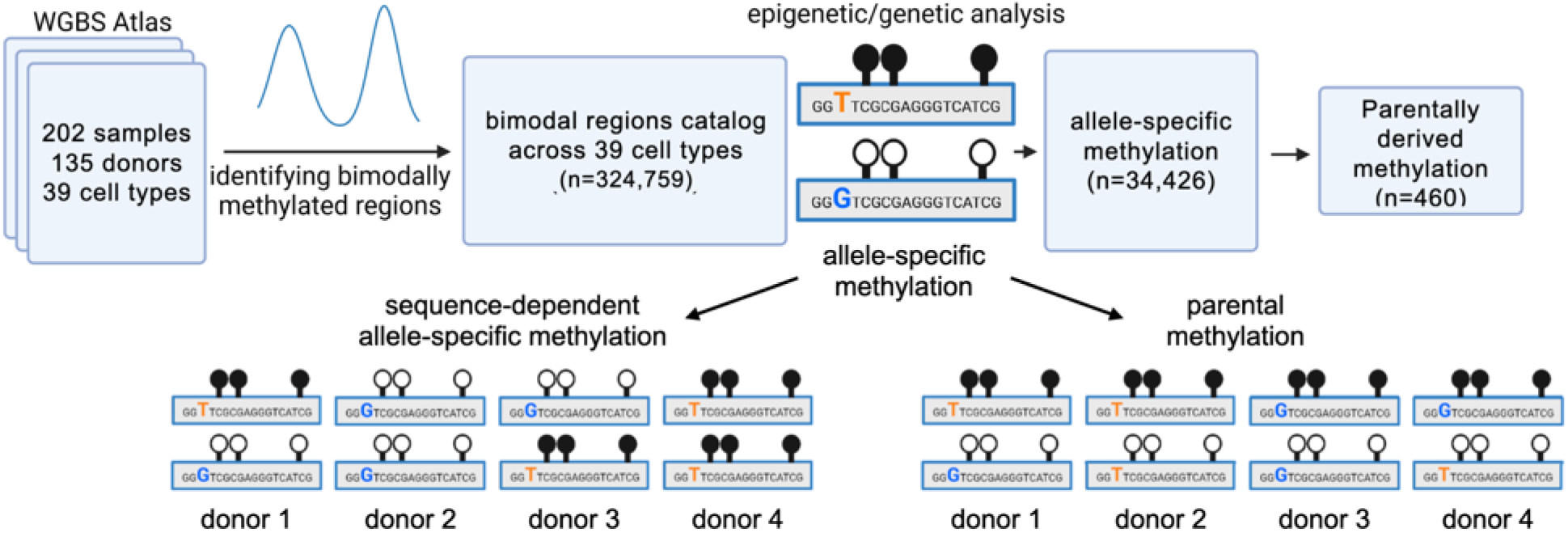
Schematic workflow: A pan-tissue human atlas of bimodal, allele-specific and parent-of-origin DNA methylation. Using fragment-level analysis, we identified 324,759 genomic loci showing a mixture of fully methylated and fully unmethylated DNA fragments. Genetic variation at neighboring SNPs was used to split the fragments by genotype, and to identify 34,426 loci that show allele-specific methylation. These were analyzed across multiple donors and classified as sequence-dependent allele-specific methylation (SD-ASMs, bottom-left) where methylation consistently segregates with one allele, or parental methylation (bottom-right) if both methylated and unmethylated epialleles exist, regardless of the genotype, suggesting that methylation is associated with parental, rather than genetic, origin. Overall, we identified 460 parental loci, including most known imprinting control regions, as well as multiple novel regions, which we associate with neighboring genes showing allele-specific expression or allelic bias. Remarkably, some of these regions also show cell-type-specific effects, including escape of allele-specific methylation.

### Identification of regions with bimodal methylation

Our initial analysis focused on identifying regions of bimodal methylation, where half of the sequenced reads are methylated, and the other half are unmethylated^32,36,37^. Such bimodality could be attributed to differential methylation in sub-populations of cells; however in the case of primary cell types purified to homogeneity from a single donor, the existence of two epialleles suggests allele-specific methylation (ASM) - either at imprinting control regions (ICRs) where one parental allele is methylated while the other is not, or at sequence-dependent ASM (SD-ASM) regions, where a heterozygous SNP is associated in cis with differential methylation^7,15^. To identify bimodal regions, we classified each DNA fragment, typically covering multiple neighboring CpG sites, as “mostly unmethylated” (U), “mostly methylated” (M), or “mixed” (X)^38^. We then calculated the percent of U/X/M fragments in each genomic position, and developed an algorithm to identify genomic regions consisting of a mixture of methylated and unmethylated fragments (Figures 2A, S1).

**Figure 2.**
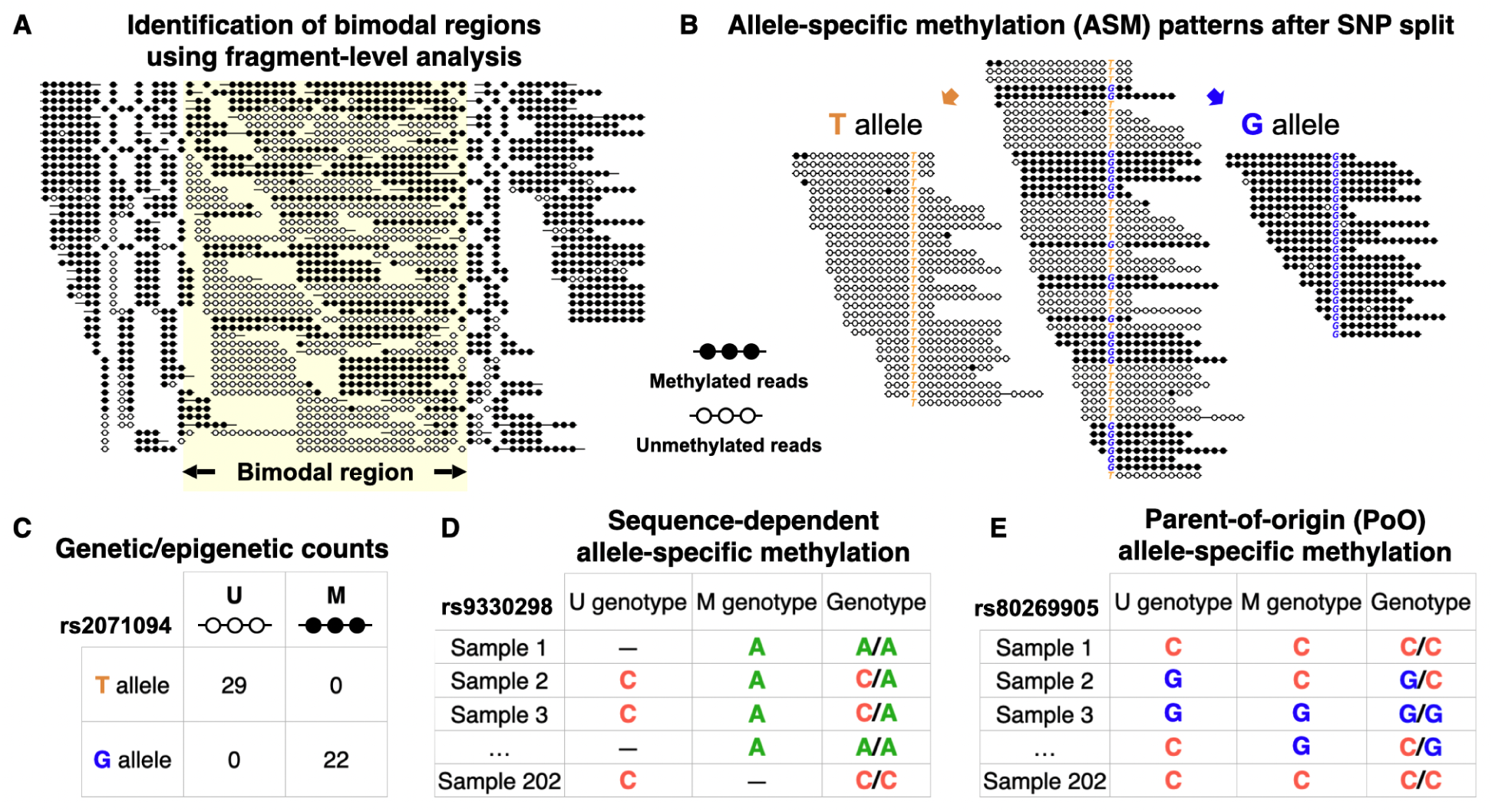
Joint genetic/epigenetic analysis across 202 samples from 135 donors identifies parental allele-specific methylation. **A)** A computational algorithm identifies bimodal regions (n=324,759), by analyzing deeply sequenced methylomes from 39 cell types. Shown is a bimodal region (chr19:54039871-54043130, hg19, highlighted) where 51% of DNA fragments are methylated (black), and 46% are unmethylated, in a colon macrophage sample purified from a single donor. **(B)** Similarly, DNA fragments from Adipocytes were split by a common T/G SNP (rs2071094, chr11:2021164) to show allele-specific methylation. Fragments carrying the T allele are unmethylated (white), whereas G allele fragments are methylated. **(C)** Contingency table of alleles by methylation, as shown in (B). All 29 unmethylated fragments are from the G genotype, whereas all 22 methylated ones carry the T genotype (adj. p-value≤7.1E-19, Fisher’s exact). **(D)** Genetic/Epigenetic table across multiple samples/cell types (rs9330298, chr1:153590254, hg19). Here, for all samples (homozygous or heterozygous), unmethylated fragments (U) have the G genotype, whereas the methylated fragments (M) are associated with the alternative T genotype, consistent with sequence-dependent allele-specific methylation (SD-ASM). **(E)** A similar table for rs80269905 (chr11:2720873), is consistent with parental imprinting. All samples are bimodal (showing both U and M fragments), and heterozygous samples are associated with allele-specific bimodal patterns but switch across different donors.

Importantly, our algorithm is based on fragment-level analysis and does not rely on fixed-sized windows, allowing for flexible and accurate determination of start and end positions, at a single base-pair resolution. Overall, we identified 324,759 regions showing a significant bimodal pattern in at least one sample (Data S1). These bimodal regions cover all (100%) known imprinting control regions, with a total of 172 Mb (5.7% of the genome, 11% of all CpG sites). On average, each individual sample shows bimodal patterns across 2.45% of CpG sites (std=1.6) or 1.15% of the genome (34.76Mb, std=0.94), with most bimodal regions (65%) showing bimodal patterns in at least 10 samples.

### Ubiquitous and cell-type-specific methylation patterns at known imprinting control regions

Most notable of these regions is the ICR for IGF2^28^ (chr11:2018812-2024740, hg19), which shows bimodal methylation patterns in all 202 samples. As Figure 3A-B demonstrates, half of the sequenced reads at this region are methylated across multiple CpGs, whereas the other half are unmethylated, consistent with bimodal allele-specific patterns originating from differential parental methylation. Similarly, the known ICRs^28^ of DIRAS3, ZDBF2/GPR1-AS, PLAGL1, PEG10, and others are ubiquitously bimodally methylated (Figure S2).

**Figure 3.**
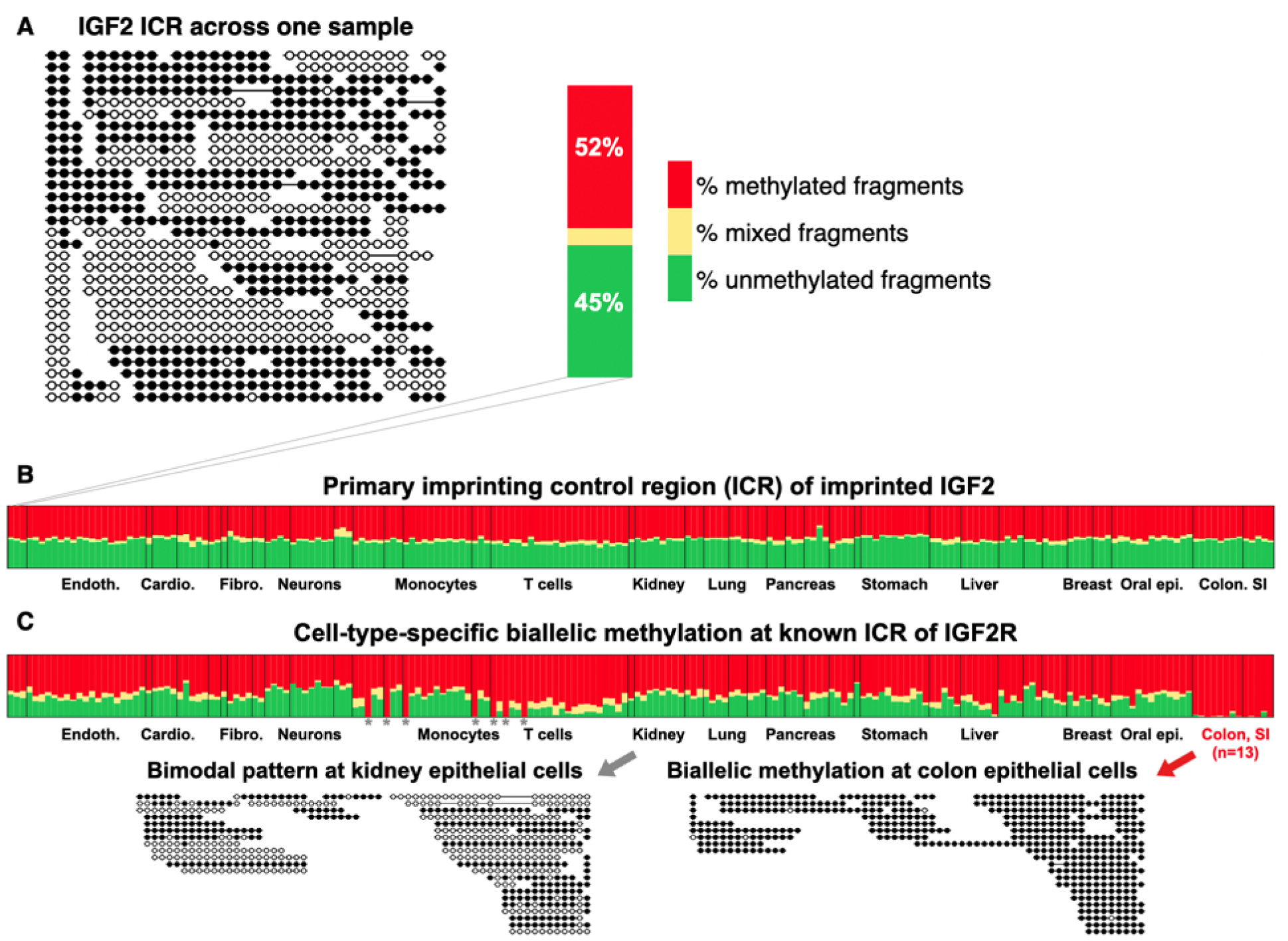
Pan-tissue analysis of bimodal methylation in known imprinting control regions (ICRs). **(A)** DNA fragments from adipocytes purified from a single donor, at the known imprinting control region (ICR) of the IGF2 gene (chr11:2018812-2024740, hg19). 52% of fragments are fully methylated (black circles) whereas 45% are fully unmethylated (white circles). **(B)** Stacked bars showing the percent of methylated (red), unmethylated (green) or mixed (yellow) DNA fragments, across 202 purified samples, spanning from 39 cell types, where a nearly balanced pattern of 1:1 ratio between unmethylated and methylated fragments is shown for this known imprinted region. **(C)** Same for the known ICR of IGF2R (chr6:160426558-160427561, hg19), showing a bimodal (imprinted) pattern across all samples, except for small intestine and colon epithelial cells (n=13 samples), where both alleles are fully methylated (right, bottom). Grey asterisks mark samples from a single donor who exhibits bi-allelic methylation.

Remarkably, not all ICRs are bimodally methylated across all adult tissues. For example, IGF2R is maternally expressed in mice, but not in humans^39^ where both alleles are expressed, purportedly due to the loss of the ncRNA Air^40,41^. Nonetheless the known ICR for IGF2R (chr6:160426558-160427561, hg19) was thought to show allele-specific methylation in all human adult tissues^40,42,43^. Using our data, we show that while the ICR is generally bimodal, it is fully methylated, across both alleles, in all 13 colon and small intestine epithelium samples (Figure 3C). Besides IGF2R, we found 13 known ICRs that show cell-type-specific alterations of the bimodal (imprinted) pattern. Intriguingly, we observed regions that became biallelically hypomethylated as well as regions that became fully methylated, suggesting high cell-type-specific plasticity at parentally methylated regions (Figures S2, S3).

Additionally, a detailed examination of known ICRs across different cell types identified fluctuations in their exact boundaries, as well as internal patches that are fully methylated in some cell types. For example, the germline ICR of H19/IGF2^28^ is bimodal in all blood samples but contains a small 700bp-long region of biallelic methylation in hepatocytes and pancreatic samples (Figure 4A-B).

**Figure 4.**
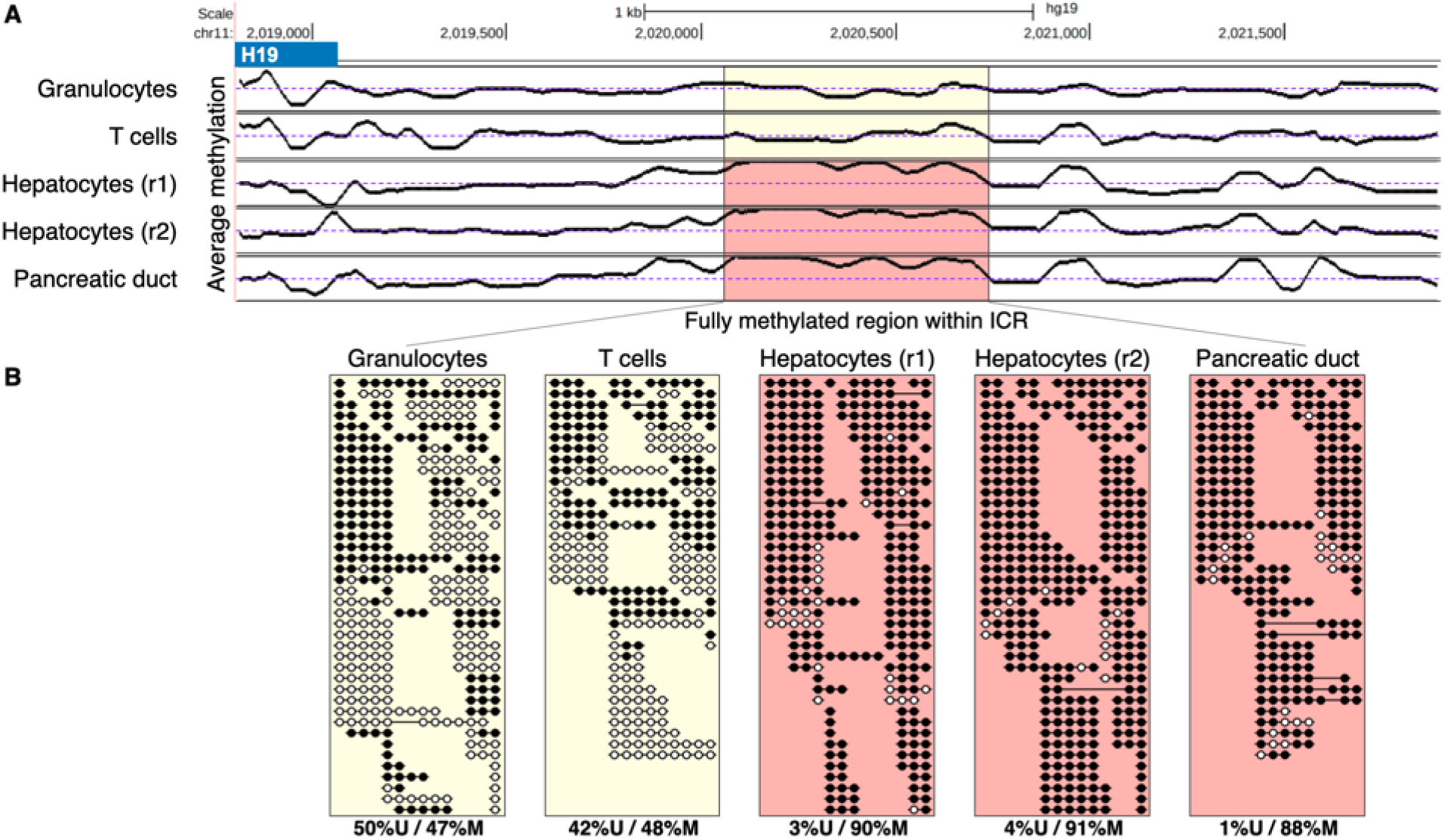
Tissue-specific loss of imprinting in intra-ICR patches. **(A)** Average CpG methylation plot (chr11:2018807-2021899) within the known ICR for H19/IGF2. The highlighted 692bp patch (chr11:2020097-2020789) shows monoallelic methylation in purified granulocytes and T cells samples (as expected) but is fully methylated in hepatocytes, and pancreas ductal epithelial samples. An average of 50% methylation is expected for known ICRs (dotted line). **(B)** Sequenced DNA fragments from within the highlighted region, revealing intra-ICR biallelic methylation. Indeed, almost all fragments from the hepatocytes and pancreas are fully methylated, compared to half of granulocyte and T cell DNA fragments.

We therefore used the genome-wide catalog of bimodal regions to automatically highlight and mask out intra-ICR regions that show a biallelic methylation pattern (hyper- or hypo-methylated), thus improving the positional definitions of known ICRs near imprinted genes (Table S1).

### Allele-specific methylation patterns segregated by SNPs

Bimodal methylation patterns within a pure cell population can originate from random patterning (metastable epialleles)^44^, from genetic polymorphisms (meQTLs), or from parent-specific mechanisms. To distinguish between these possibilities, we developed a statistical procedure to test whether epialleles are associated with specific SNPs. We examined gnomAD^45^ and identified common SNPs (minor allele frequency, MAF >1%) that intersect with our set of 325k bimodal regions. In 152k regions, at least one sample exhibited both bimodal methylation and heterozygous SNP (Figure 2B-C). Overall, we identified 55,271 SNPs that segregate within 34,426 unique allele-specific methylation regions (Table S2). These regions show a bimodal methylation pattern across DNA fragments that cover ≥3 CpG sites, which is segregated in at least one sample. Thus, 4% of SNPs tested were found to associate with ASM, which is more conservative than previous estimates^4,7,9^. Note that for the majority of bimodal regions, where no informative SNP were found, we are unable to assess if their methylation is sequence-dependent or not. Since sequence reads are typically ∼200bp long, it is impossible to assess the presence of distant genetic variants controlling methylation. Thus, the 34k regions that show bimodal methylation associated with SNPs is a lower bound, and the actual magnitude of sequence- or parental-controlled methylation is likely much larger.

### Parental or sequence-dependent allele-specific methylation?

Once regions showing allele-specific methylation were identified, we examined their genetic and epigenetic patterns across multiple samples. As figure 2D demonstrates, regions of sequence-dependent allele-specific methylation (SD-ASM)^7,15,16^ show similar associations between allele and epiallele across donors, including biallelic methylation patterns for homozygous donors. Conversely, parental methylation will show bimodal patterns across multiple donors, regardless of genotyping (Figure 2E). Using stringent statistical thresholds, we identified 460 putative parental regions, covering most known imprinting control regions (45/55, 82%)^27,28^. The remaining parental regions we identified include 78 regions adjacent to known imprinted genes (up to 100Kb), 14 that reside near known ICRs (≤100Kb), and 373 parental regions whose function is yet to be determined, of which 347 are novel (Table S3). Figure S4 shows the distribution of bimodality, parental methylation, and known ICRs across the human genome (hg19), and figure S5 shows the distribution of the number of samples exhibiting bimodality, per parentally methylated region.

### Validation of parent-of-origin methylation at novel tissue-specific locus

To validate tissue-specific parent-of-origin methylation patterns, we selected one novel parental region and studied its methylation patterns across different tissues and cell types. We focused on a genomic region that is fully methylated in blood but bimodal in epithelial cells (Figure 5A), and performed a targeted-PCR methylation sequencing across 33 mother-father-child trios (Tables S9, S10). This allowed us to capture the epigenetic landscape across multiple CpGs while genotyping the target SNP at each sequenced molecule. As predicted by the WGBS atlas data, unmethylated fragments in this locus were associated with the paternal allele in all discernible cases (Figure 5B-F).

**Figure 5.**
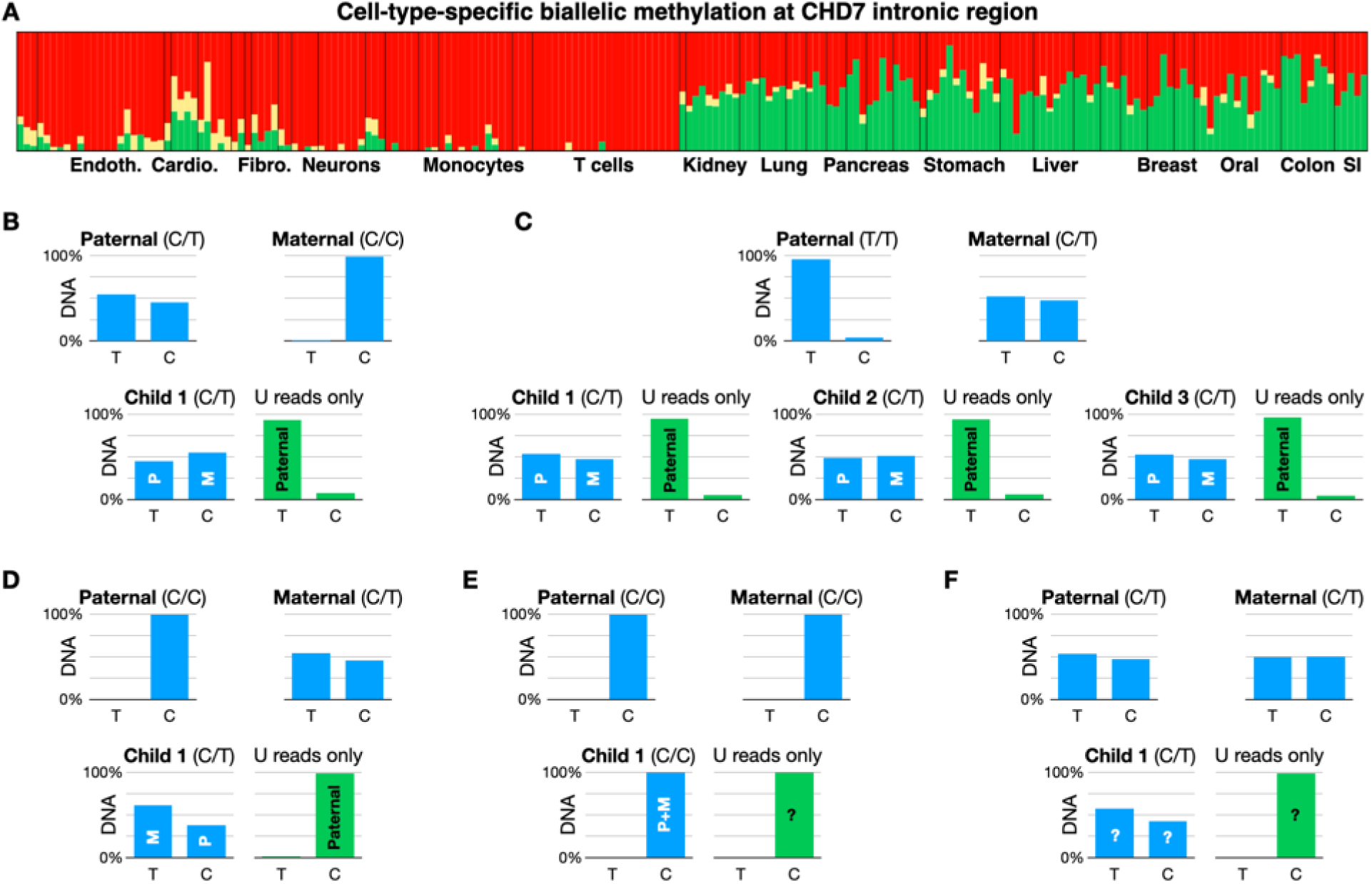
Validation of a novel cell-type-specific parentally methylated region. **(A)** Fragment-level analysis of a novel parent-of-origin cell-type-specific region we identified (chr8:61627212-61627412) shows biallelic methylation (red) in endothelial cells, neurons, fibroblasts, and blood cells, but bimodal methylation patterns in hepatocytes and epithelial cells (1:1 ratio of fully methylated and fully unmethylated sequenced fragments of ≥3 CpGs). **(B-F)** Genetic/epigenetic analysis of parental methylation in tongue epithelial cells, validated across 15 families (a total of 33 children and their parents). For each trio, we used targeted-PCR next-generation sequencing (after bisulfite conversion) to measure the genotype (rs7826035, C/T, chr8:61627312) and the methylation status of six CpG sites (chr8:61627190-61627349, measured on the bottom strand). **(B)** A trio (family ID IMP017) showing homozygous C/C for the mother, with C/T heterozygosity for the father. The child T allele is therefore of paternal origin. Blue bars correspond to relative allelic read count. Unmethylated DNA fragments (green bars) are limited to the parental T allele, suggesting maternal-specific methylation. **(C)** same as (B) for a family with three heterozygous children (IMP012). **(D)** A family where the T allele of heterozygous child 1 is maternal, whereas unmethylated fragments (green) are all from the paternal C allele. Two additional siblings are C/C homozygous and not shown (family ID IMP005). **(E-F)** Examples of a C/C homozygous family and a C/T heterozygous family, where the parent-of-origin of unmethylated fragments cannot be associated with a parent-of-origin (family IDs IMP014, IMP011). All remaining families were homozygous (C/C, not shown).

### Parental allele-specific methylation at regulatory regions

Parent-of-origin differential methylation is key to regulating allele-specific expression of imprinted genes. We therefore used functional annotations to test whether our catalog of parentally methylated regions is enriched for various genomic features. Indeed, we have observed significant local enrichment for promoters (68% of regions, chromHMM annotations), enhancers (8%)^46^, transcription factor binding sites^47^ (Tables S4, S5, Figure S6), histone marks of active gene regulation (H3K27ac, 56%)^48^, and origins of replication (46% of parental regions)^49^.

These observations highlight the regulatory role of the parental regions we identified through various molecular mechanisms. Furthermore, gene-set enrichment analysis for the putative parental regions we identified showed a 47-fold enrichment for “maternal imprinting” (FDR ≤ 2.6E-42)^50^. Intriguingly, over 46% of parental regions (213/460) overlap with origins of replication^49^, compared to 17% expected by random (FDR ≤ 9E-58), with an average S/G1 ratio of 1.46 at parental regions (Figure 6D).

**Figure 6.**
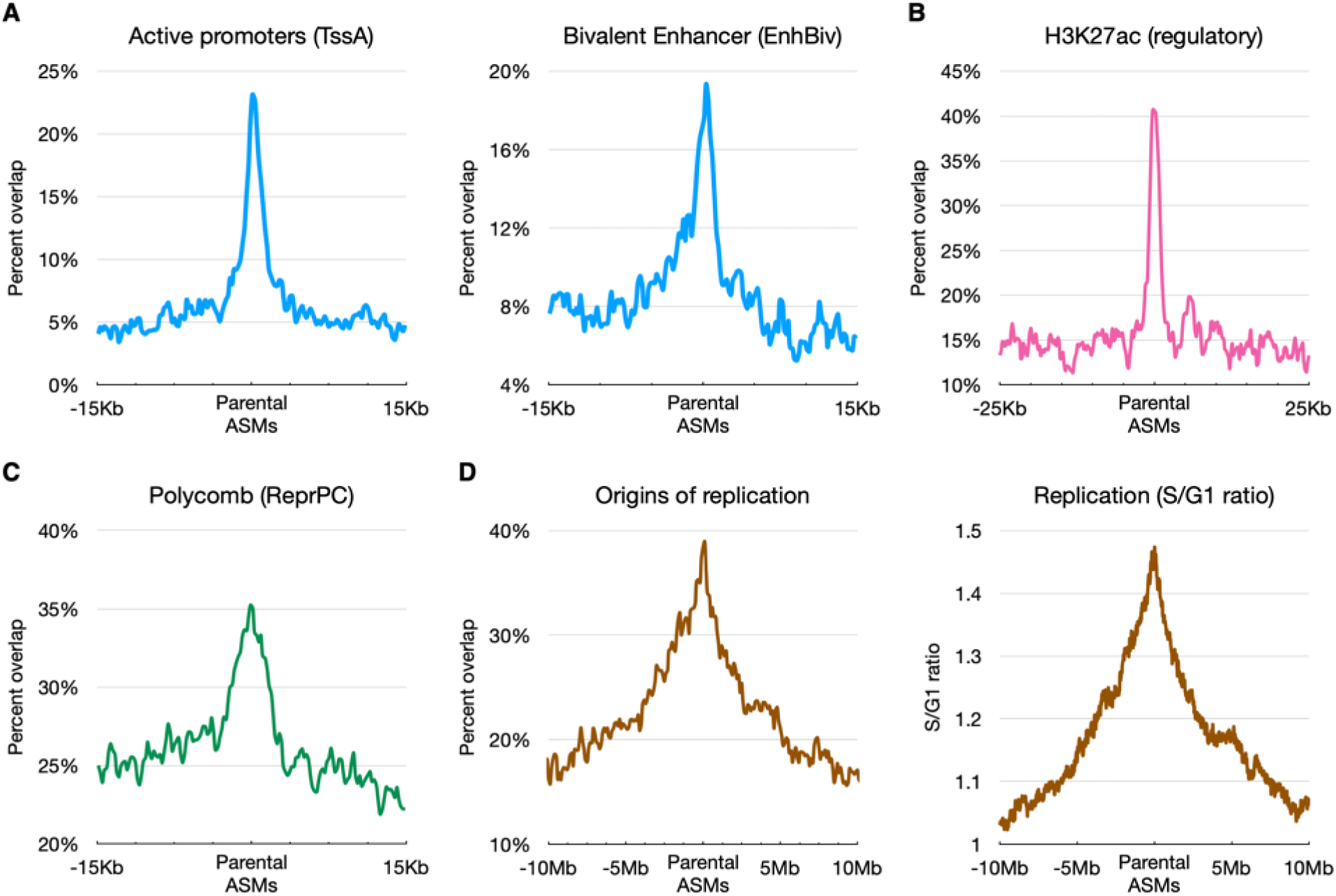
Enrichment of functional annotations at putative parent-of-origin allele-specific regions. **(A)** Percent of parentally-methylated ASMs annotated as active promoters and bivalent enhancers (chromHMM TssA and EnhBiv terms, respectively). **(B)** Enrichment of gene regulatory activity, based on H3K27ac peak annotation in 387 ChIP-seq experiments (AREs)^49^. **(C)** Enrichment for Polycomb repressive regions (chromHMM ReprPC). **(D)** Local enrichment for origins of replication, identified using peaks of nascent strand DNA (left), as well as early replication, measured as the ratio of S to G1 phase DNA fragments^49^.

### Parental methylation near imprinted and allele-biased genes

Having identified regions exhibiting putative parent-of-origin allele-specific methylation, we sought to associate them with imprinted genes and their ICRs (Table S6). Combined with Data S1, this map represents a comprehensive catalog of imprinting control regions and parental DMRs in humans. Remarkably, we identified novel parentally methylated regions near seven imprinted genes for which no ICR was previously found, including PAX8/PAX8-AS1, GNG7, ZNF215, UTS2, AXL, and KIF25 (Table S6). As figure S7 shows, the novel region we identified 18Kb upstream to PAX8 (chr2:113953706-113955952) shows bimodal methylation in neuronal cells, on par with allele-biased gene expression in the brain^16^.

We therefore used expression data from GTEx to examine allele-specific expression across the human genome, using 15,253 samples collected from 838 donors^51,52^, and identified 2,246 genes exhibiting a significant bias in at least one tissue type. Of these, 216 genes are located near parental/bimodal regions in matched cell types (≤250Kb), compared to 111 genes expected at random (std=10; permutation test p≤4E-34, Table S7). These findings further support the idea that bimodally methylated regions control allele-specific gene expression in *cis*.

### Putative mechanism underlying tissue-specific biallelic expression of imprinted genes

Despite the common canonical examples wherein imprinted genes are monoallelically expressed in all cell types, in certain instances imprinted genes were shown to “escape” parental repression and to exhibit biallelic expression in a tissue-specific manner^16,53,54^. One notable example is IGF2, which is monoallelically expressed in most tissues, but is expressed in the liver from both maternal and paternal alleles^16^ (Figure 7A). The mechanisms underlying the parental regulation of IGF2 are well studied - in the paternal allele, the primary ICR is methylated to prevent binding of the insulation factor CTCF, thus allowing IGF2 activation by distal enhancers. Conversely, the maternal allele is unmethylated and CTCF is bound, leading to distal activation of H19, but not IGF2^53,55^ (Figure 7B). However, the regulatory mechanisms underlying biallelic expression of IGF2 in the liver have not been elucidated^53,55^.

**Figure 7.**
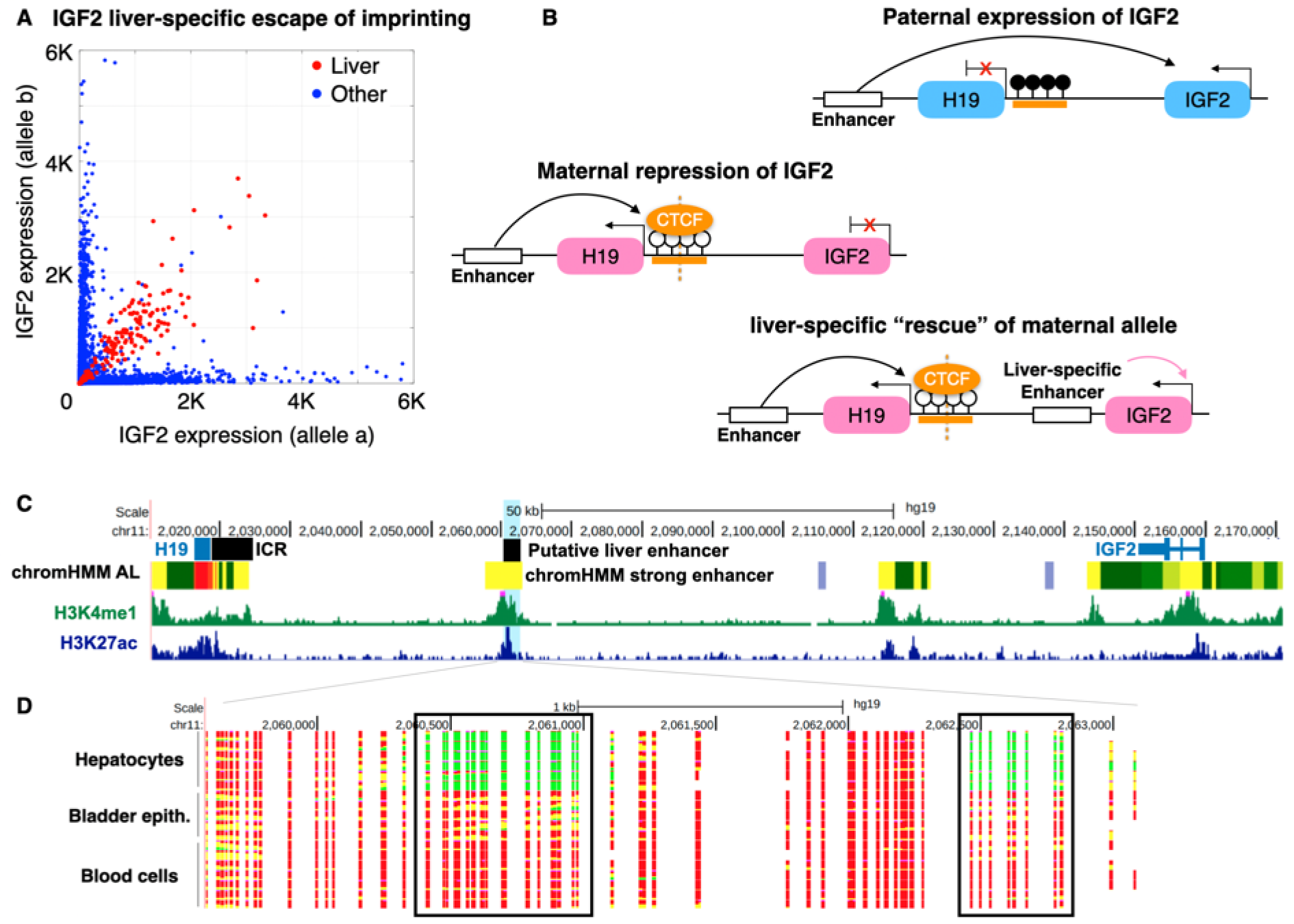
Putative novel enhancers overcome IGF2 imprinting in human hepatocytes. **(A)** Allele-specific read counts for IGF2 (GTEx) show biallelic expression of either the A (X-axis) or B (Y-axis) alleles (blue dots). Conversely, mRNA from liver cells (red) show a diagonal, 1:1 allelic ratio, consistent with liver-specific escape of imprinting. **(B)** Known imprinting control mechanism for IGF2/H19. In the paternal allele (top), methylated CpGs (black lollipops) prevent CTCF from binding the ICR, facilitating the activation of IGF2 by a distal enhancer. Conversely, in the maternal allele CTCF binds the unmethylated ICR, acting as an insulator^53,55^. We propose a mechanism by which liver-specific enhancers activate IGF2 in both maternal and parental alleles, thus escaping maternal imprinting. **(C)** Genomic view of the H19/IGF2 locus. The putative liver enhancers (highlighted in blue) show strong H3K4me1 and H3K27ac ChIP-seq peaks in adult liver tissue (green and blue tracks), but not elsewhere (not shown). chromHMM adult liver (AL) track shows a putative annotation of this region as active enhancers (yellow). **(D)** Fragment-level analysis using our whole-genome methylation atlas shows fully unmethylated fragments (green) in six hepatocyte samples, at two adjacent putative enhancers (black frame), compared to fully methylated fragments in other cell types, where IGF2 is maternally imprinted.

Our data shows bimodal methylation at the IGF2 ICR across all samples (Figure 3A-B) including hepatocytes, suggesting that a different mechanism is underlying liver-specific maternal activation. We identified two genomic regions in the vicinity of IGF2 that are fully unmethylated in hepatocytes (in both alleles) but are fully methylated elsewhere in the human body^3^. These two putative enhancers are also characterized by enhancer-specific chromatin marks (H3K4me1 and H3K27ac), and were annotated as putative liver enhancers by chromHMM^56,57^.

These findings suggest the presence of liver-specific enhancers that activate IGF2 specifically in the liver, including in the otherwise silenced maternal allele, thus overriding ICR-driven maternal allele repression (Figure 7B-D). Based on this example, we developed a computational score to compare allele-specific gene expression data^52^ with the presence of differentially unmethylated neighboring regions, and identified putative cell-type-specific enhancers for 34 imprinted genes (Table S8), suggesting a general mechanism for tissue-specific escape from parental imprinting.

## Discussion

We describe here a comprehensive atlas of allele-specific DNA methylation in all major human cell types, based on deep whole-genome bisulfite sequencing of DNA from freshly isolated cells. A unique strength of this atlas is that it is based on DNA from purified cell types, allowing identification of bimodal methylation patterns that are due to within-cell-type phenomena, rather than cell mixture effects.

Overall, we identified 325k genomic loci that exhibit bimodal methylation patterns in at least one sample, covering 5.7% of the genome and 11% of CpGs (average of 2.45% of CpGs in bimodal regions, per sample). We observed differences in bimodality both between and within cell types, due to cell-type-specific effects in parental methylation, as well as genetic differences between individuals in regions associated with sequence-dependent allele-specific methylation. In 10% of bimodal regions (34k loci), we were able to identify SNPs that segregate with methylation, demonstrating allele-specific methylation. The remaining loci may feature allele-specific methylation with distant sequence determinants that cannot be captured with short-read sequencing, or cases in which one random allele per cell is methylated, similarly to the situation in the mammalian X chromosome^14^. We were able to identify some definitive examples of the latter (i.e. bimodal methylation that is neither parental nor allele-specific), therefore the scope of this phenomenon remains to be determined. Additionally it is possible that in some of our purified samples there are hidden sub-types of cells that harbor distinct methylation patterns (e.g. different types of pancreatic beta cells, all expressing insulin).

The comprehensive catalog of allele-specific and parental methylation presented here is consistent with previously published maps of parentally-derived ASM^7,15,30,31^. The main differences we present stem from our use of homogenous samples of various purified cell types, deeply sequenced across multiple donors and cell types. Two previous studies utilized samples with uniparental disomy and conducted bisulfite sequencing from blood to collectively discover 92 parentally methylated regions^30,31^. Of the 36 regions identified by Court et al., we classified all 36 as bimodal, with 32 supported by SNPs indicating parental methylation. Joshi et al. identified 79 regions, of which we classified 76 as bimodal, with 44 supported by heterozygous SNPs in our samples. A more recent study utilized a WGBS blood dataset consisting of 285 samples with complete parent-of-origin genetic information for over 1.9 million SNPs^15^, and identified regions of parental ASM. 76% of these (174/229) were recognized as bimodal in our atlas, and 85 regions had enough samples with heterozygous SNPs to be classified as parentally methylated. The remaining 55 regions did not pass our strict threshold for statistical significance for bimodality. Thus our atlas reproduces most regions identified by previous studies in blood cell types, and highlights the importance of studying allele-specific methylation in a variety of cell types.

The association of methylation patterns with SNPs allowed us to identify 460 genomic regions with putative parental methylation. Reassuringly, most known imprinting control regions - loci with parentally regulated methylation - are present in this list (45/55). In addition, 78 of the loci we discovered are associated with known imprinted genes, providing a putative mechanism of regulation, and 14 loci reside in vicinity to known ICRs (up to 100kb). The remaining 373 loci represent a comprehensive landscape of parental allele-specific methylation. We validated one such locus, showing parentally-associated allelic differences in epithelial - but not blood - cells, using trio analysis in swab samples. The imprinting status of the other loci as well as their function merit further investigation. Further, the validation of this methodology and the existence of tongue epithelium-specific parental ASM markers exemplifies the plausibility of using tongue swabs to detect congenital imprinting-related disease. To date, only screens using blood samples have been developed^27^.

The substantial number of loci with putative parental allele-specific methylation allowed us to characterize these regions, revealing enrichment for regulatory regions (bearing chromatin marks of enhancers and promoters) and for polycomb targets, consistent with a role in regulation of monoallelic expression of nearby genes. Putative imprinted loci also tend to reside near origins of DNA replication, raising the testable hypothesis that parent-of-origin dependent asynchronous DNA replication controls parent-of-origin-dependent allele-specific methylation^58,59^. Further analysis is required to investigate the relationship between cell-type-specific allele-specific methylation and cell-type-specific asynchronous DNA replication.

One striking phenomenon emerging from the atlas is tissue-specific escape from imprinting. Previous studies of imprinting mostly focused on blood cells, and assumed that parental imprinting in the gametes persists in all tissues^19–22^. The presence of multiple rarely studied cell types in the atlas exposed the fact that many parentally determined loci (including almost a quarter of known ICRs, 13/55) escape imprinting and become fully methylated or fully unmethylated in specific cell types. The biological significance of this fascinating phenomenon and the underlying molecular mechanisms likely vary, depending on the affected gene and cell types. Our analysis of select cases suggests one way by which specific cell types may overcome monoallelic expression imposed by parent-of-origin-dependent methylation. As we showed for IGF2, the presence of a tissue-specific enhancer near the gene (in this case, a proximal enhancer that is fully unmethylated only in hepatocytes) likely allows expression from both alleles, despite monoallelic availability of the remote enhancer.

The comprehensive atlas of parentally-imprinted and sequence-dependent DNA methylation in a variety of human cell types provides a platform for additional computational and wet lab analysis, to address the fundamental question of how, why, and to which extent cells distinguish between different alleles of the same gene, a phenomenon with important biological and clinical implications.

## Methods

### Materials Availability

All WGBS sequenced and derived data is available on GEO (accession no. <u>GSE186458</u>). Targeted PCR sequencing of trios at the novel tissue-specific region will be made available by the date of publication. The code will be made available by the date of publication. Any additional information required to reanalyze the data reported in this paper is available from the lead contact upon reasonable request.

### Experimental model and study participant details

Sequencing data (150bp paired-end reads, 984 million pairs per sample) for the atlas were mapped to the human genome (hg19) and analyzed using the software suite wgbstools^38^ (github.com/nloyfer/wgbs_tools) as described in Loyfer et al^3^.

The clinical study of the trios was approved by the ethics committee of the Hadassah Medical Center. Procedures were performed in accordance with the Declaration of Helsinki (HMO-0198-14). Swab sampling donors have provided written informed consent. Participant details are provided in Table S9.

This study was approved by the Hadassah Hospital Institutional Review Board and all participants provided signed informed consent (adults) or parental consent (children).

### DNA processing

Swab samples were collected with an inoculating loop by swabbing it against the tongue of healthy donors for 20 seconds and breaking the inoculating loop into a 2ml Eppendorf tube with 200ul of PBS. The sample was saved in a freezer at −20C until extraction. DNA extraction was performed via DNeasy Blood and tissue kit (QIAGEN) according to the manufacturer’s instructions with the following change: incubation time of AL buffer was performed overnight. The DNA concentration was measured using Qubit High Sensitivity double-strand molecular probes (Invitrogen) and bisulfite treatment (Zymo). Bisulfite-treated DNA was PCR amplified using primers (Table S10) specific for bisulfite-treated DNA but independent of methylation status at monitored CpG sites or genotype. We used a multiplex 2-step PCR protocol as described in Neiman et al^63^. Pooled PCR products were subjected to multiplex NGS using the NextSeq 500/550v2 Reagent Kit (Illumina). Sequenced reads were separated by barcode, aligned to the target sequence, and analyzed using custom scripts written and implemented in R. Reads were quality filtered based on Illumina quality scores. Reads were identified by having at least 80% similarity to target sequences and containing all the expected CpGs in the sequence. CpGs were considered methylated if “C” was read at a CpG site and were considered unmethylated if “T” was read. The efficiency of bisulfite conversion was assessed by analyzing the methylation of non-CpG cytosines. We then determined the fraction of molecules in which all CpG sites were unmethylated. Further ASM analysis was similar to those of the WGBS samples. Reads were segregated by genotype at SNP rs7826035 (C/T, chr8:61627312l). For C/T SNPs only bottom stranded reads are considered. Tongue swab samples contain a mix of blood and tongue epithelial cells. Since the novel region detected is bi-allelically methylated in blood cells we verified that unmethylated fragments (which must come from tongue-epithelial cells) always contained the paternal genotype. All samples were required to contain at least 1000 sequenced fragments and each C/T allele was determined to be present if at least 40% of reads exhibited the relevant genotype. We classified children as exhibiting paternal-specific demethylation if over 95% of unmethylated fragments contained the paternal genotype.

### Data and identification of bimodal methylation regions

Sequenced read pairs were then merged, and classified as hyper-methylated (M) if covering three CpGs or more, with an average methylation (per fragment) ≥ 65%. Similarly, hypo-methylated (U) fragments were defined as having average methylation ≤ 35%. Fragments with less than 3 CpG sites are ignored, and the remaining fragments, with an avg. methylation of (35%-65%), were classified as mixed (X). For each CpG site, we calculated the {U,X,M} proportions across all overlapping fragments with ≥3 CpGs. Bimodal regions were then defined as contiguous regions (≥ 5 CpGs) where the proportion of both hyper- and hypo-methylated fragments (i.e., U, M) is ≥ 20%.

We then devised a statistical test to distinguish between a null hypothesis model H_0_ of one epi-allele (possibly showing ∼50% methylation, on average), vs. a mixture model H_1_ of two equally likely epi-alleles (A, B), corresponding to DNA fragments originating from the methylated and the unmethylated alleles. Both models assume conditional independence between neighboring CpGs, and model the expected methylation at the i’th CpG (for a given epiallele) using a Bernoulli parameter *P*(*r*_*i*_|*epiallele*) where *r*_*i*_ is an indicator for whether the DNA fragment r shows that CpG i is methylated. For H_0_, these probabilities could be estimated using a maximum likelihood estimator, based on the empirical probability of methylation at the i’th CpG (beta value). For H1, we used the expectation-maximization (EM) algorithm, to iteratively infer the posterior probability that each fragment r is associated with each epiallele *Pr*(*A*|*r*), *Pr*(*B*|*r*), and estimated the expected methylation *P*(*r*_*i*_|*A*), *P*(*r*_*i*_|*B*) of each CpG i given the two epialleles.

In H_0_ we define the probability to be methylated at CpG i as: θ_*i*_.

Thus, the likelihood of one region, based on H_0_, could be viewed as the product of likelihood across all fragments:

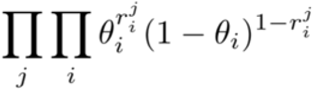

Alternatively, based on H_1_ where the probability to be methylated on allele A is ^1^ and on allele B is θ ^2^, the likelihood could be described as:

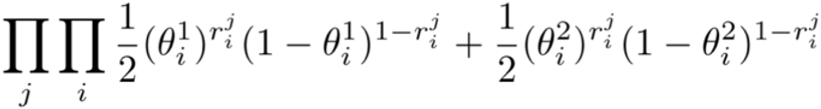

We infer the posterior probability *Pr*(*A*|*r*) per read. We assign each read to an allele by choosing the allele with the maximum posterior probability (hard assignment). We then calculate the expected methylation probability per CpG, for each of the A or B epialleles (expected counts).

Finally, we applied a log-likelihood ratio test to estimate the statistical significance of each bimodal region (comparing the two-epiallele mixture model H_1_ with the nested, single-epiallele, H_0_ model), using the software package wgbstools (test_bimodal function)^38^. p-values were then corrected for multiple hypotheses using the Benjamini-Hochberg FDR correction scheme^64^.

To further extend identified bimodal regions beyond domains of densely located CpG (resulting with sequenced fragments covering ≥ 3 CpGs), with allow expansion into flanking regions, as bimodality is maintained (namely, while both hypo- and hyper-methylated fragments are ≥ 20%).

This computational procedure was applied to each sample independently, to account for genetic and environmental changes. We then set the start and end position of each bimodal region to the closest methylation block boundary, as determined using a genome-wide segmentation of the genome, using the wgbstools package as described in Loyfer et al^3,38^.

### Allele-specific methylation

To associate bimodal regions with two independent allele-specific methylation (ASM) patterns, we integrated these regions with 1,360,985 SNPs showing a minor allele frequency (MAF) ≥ 1%, using GnomAD^45^ data (Ashkenazi Jewish population). For each such SNP, we retained the heterozygous samples (≥ 5 fragments from each allele), and built a contingency table comparing the number of U/X/M fragments (≥3 CpGs) from each individual genotype. Fisher’s exact test was then used to test for association between allele and methylation patterns, followed by Benjamini-Hochberg FDR correction^64^ (Table S2).

### Sequence-dependent and parent-of-origins ASMs

Once allele-specific methylation (ASM) was identified, per sample, we performed a broad cross-sample analysis to test for sequence-dependent (SD-ASM) and parent-of-origin effects. For SD-ASM, all samples (of a given cell type, or in general) are expected to show concordance between genotype and methylation. That is, regardless of donor heterozygosity, all fragments from a given allele (genotype) are expected to be methylated, whereas fragments from the alternative allele not. Conversely, parent-of-origin effects are expected to show bimodality regardless of heterozygosity, and switch alleles between unrelated donors. Specifically, we require ≥3 samples to exhibit ASM and that the association between genotype and hyper/hypo methylation switch across samples. Thus if one sample allele A is associated with unmethylated epialleles and in another sample allele A is associated with methylated epialleles, the region is classified as putatively parent-or-origin derived ASM, as opposed to SD-ASM. Parent-of-origin ASMs were defined as novel if they were not identified in the previous studies of Zink et al., Court et al., Joshi et al., and Orjuela et al^15,29–31^.

### Gene enrichment analysis

These putative parent-of-origin ASM regions were compared against all imprinted genes, as reported by Tucci et al^65^, as well as known imprinting control regions (ICRs) from Monk et al^28^. For the definition of promoters, bivalent enhancers, and polycomb repressive we used chromHMM genomic annotations^66^. Each type was merged (“bedtools merge”) across all cell types. Active regulatory elements were determined using H3K27ac ChIP-seq data^48^. Origins of replication and S/G1 ratio bedGraph files were taken from Mukhopadhyay et al^49^. To test enrichment, we measured the intersection of the merged annotation track (bigwig) with the list of putative parental-ASM regions (“bedtools intersect”). Statistical significance was estimated using a permutation test of 100 random length-preserving chromosome-wide shuffles, fitted using a Normal distribution.

### Identifying biased allelic expression

Allele-specific expression data from GTEx (https://gtexportal.org/home/datasets, v8 phASER haplotype matrix) was used. For each gene, a background noise model was applied by fitting a Normal distribution for the A vs. B allele-specific read count, at discrete bins of expression levels. Any gene/sample with allelic bias ≥ 3 standard deviations was classified as showing allelic bias.

### Associating tissue-specific allelic bias with bimodal methylation

Using the allele-specific read count data from GTEx, we classified a tissue/gene pair as showing allelic bias if ≥25% of samples in that tissue show allelic bias. Cell-type-specific bimodality was defined as genomic regions for which ≥90% of sequenced samples are classified as bimodal. We identified 2,246 such genes. GTEx tissues and our purified cell types were matched across 23 tissues/cell types, and the mutual information across genes with allelic expression and bimodal regions (≤250Kb away) was computed. We compared the number of genes identified to what happens at random using permutation testing. We selected 2,246 genes at random and counted the number which were near (<=250kb away) parentally methylated regions with bimodal patterns in matching cell types as those exhibiting bias. We ran 50 iterations of random permutations.

### Escape from imprinted expression mechanisms

For every imprinted gene we find those which exhibit biallelic expression in at least one tissue type. Each gene/sample is classified as biallelically expressed if the allelic bias is within one standard deviation of the noise model described above. Tissue/gene pairs are classified as biallelically expressed if at least 25% of samples show biallelic expression. For each such gene/tissue we search for hypomethylated DMRs using wgbstools’ find_markers command with the following parameters: “--min_cpg 5 --delta_quants 0.35 --tg_quant 0.15 --bg_quant 0.4”. For our analysis of IGF2 (Fig. 6), ChIP-seq from Roadmap Epigenomics’ Adult Liver was used (H3K27ac and H3K4me1 from Donor 3, DNA_Lib 1057)^56^, as well as chromHMM primary annotations for Adult Liver^57^.

### Quantification and statistical analysis

Quantification of bimodal regions, ASM, and putative parent-of-origin methylation are described above. Bimodal regions are detected using the EM algorithm to fit model parameters and then a log odds ratio test is used to determine significance (described above). FDR is used to correct for multiple hypothesis testing, with a threshold of 0.05. Fisher’s exact test is used to determine significance of ASM with an FDR threshold of 0.01. All regions and their significance can be found in Data S1 and Table S2. Enrichment analysis used permutation testing with a FDR threshold of 0.05, as described above.

## Supplemental Information

Data S1. Genome-wide list of bimodal methylation regions by sample

Table S1. Known ICR coordinates, with regions of biallelic methylation masked Table S2. List of SNPs associated with cell-type-specific ASMs

Table S3. Genome-wide list of SNPs associated with parental ASMs Table S4. Transcription factor enrichment analysis (ReMap)

Table S5. Chromatin states enrichment analysis (chromHMM)

Table S6. Genes exhibiting allele-specific expression, with parental ASM

Table S7. Genes exhibiting escape-from-imprinting, with tissue-specific DMRs

Table S8. Genes exhibiting tissue-specific imprinting, with putative parentally methylated ASMs Table S9. Novel parental-ASM participant information

Table S10. Novel parental-ASM locus and primer information

## Acknowledgements

We thank Howard Cedar, Sagiv Shifman, Itamar Simon, Agnes Klochendler, Nir Friedman, and members of the Kaplan and Dor labs for insightful discussions. This research was supported by research grants from GRAIL (to T.K, Y.D, and B.G), the Center for Interdisciplinary Data Science Research (to T.K., Y.D. and B.G.), The Israel Science Foundation (grants 1250/18 and 259/23 to T.K.), and by the Ministry of Innovation, Science & Technology, Israel (grant 0005421).

## Author Contributions

Conceptualization, JR, RS, BG, YD, TK; investigation, JR, AP, JM, NL; writing and editing, JR, YD, TK; review & editing, all authors; supervision, YD, TK; funding, YD, TK.

## Extended data figures

**Supplemental Figure 1.**
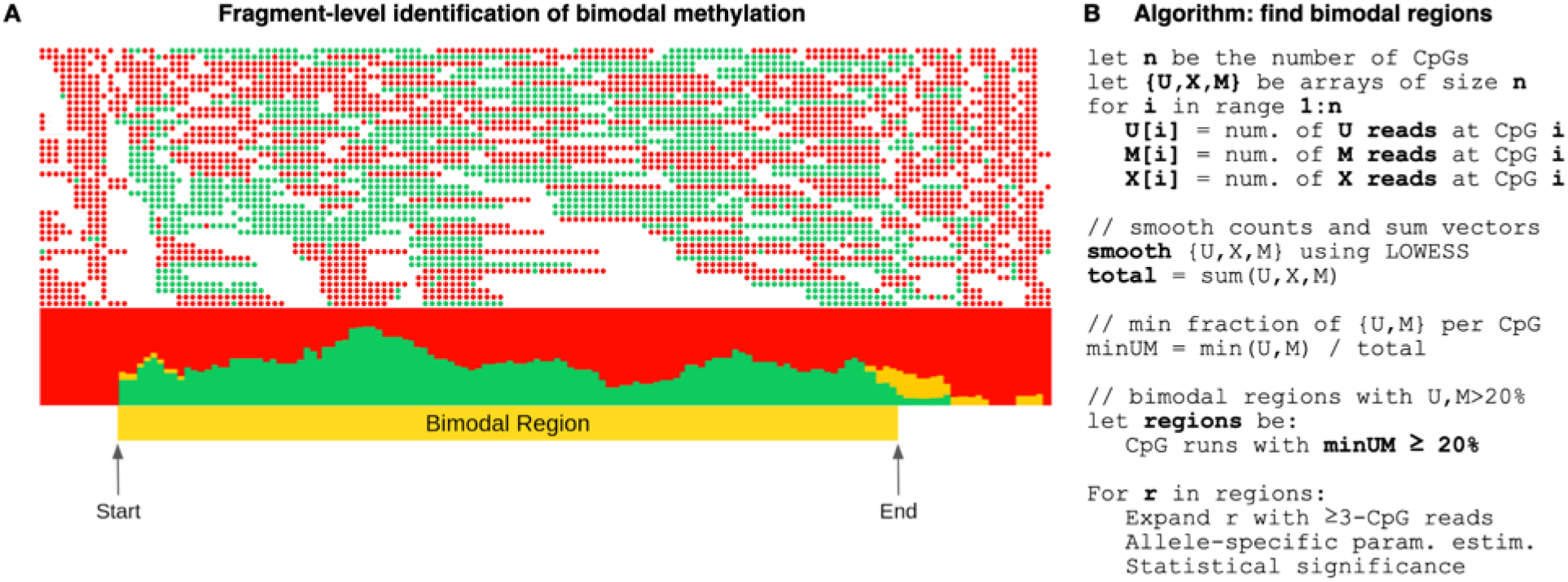
Algorithm for the identification of bimodal regions. **(A)** every read is separated by a white space. Reads are classified as either **U** (less than 35% methylated CpGs per fragment), **X** (not **U** or **M**), or **M** (more than 65% methylated CpGs per fragment). The figure below shows the cumulative proportion of U/X/M reads across each CpG, with arrows marking where the min(U,M) fragment proportion is above 20%. We classify a region as bimodal if the proportion of both **U** and **M** reads is ≥ 20% across neighboring CpGs and passes the statistical test. (**B**) Pseudo code of the efficient linear-time algorithm which identifies the bimodal regions. The algorithm works as visualized in **A**. The algorithm begins by transforming a WGBS bam file into a 3-dimensional array whereby each entry of the array represents a CpG site. At each index of the array we store how many *U/X/M* reads intersect this CpG site. We apply LOWESS smoothing on each array individually and we find those regions of contiguous CpG site where the proportion of **U** and **M** reads intersecting that site are at least 20%.

**Supplemental Figure 2.**
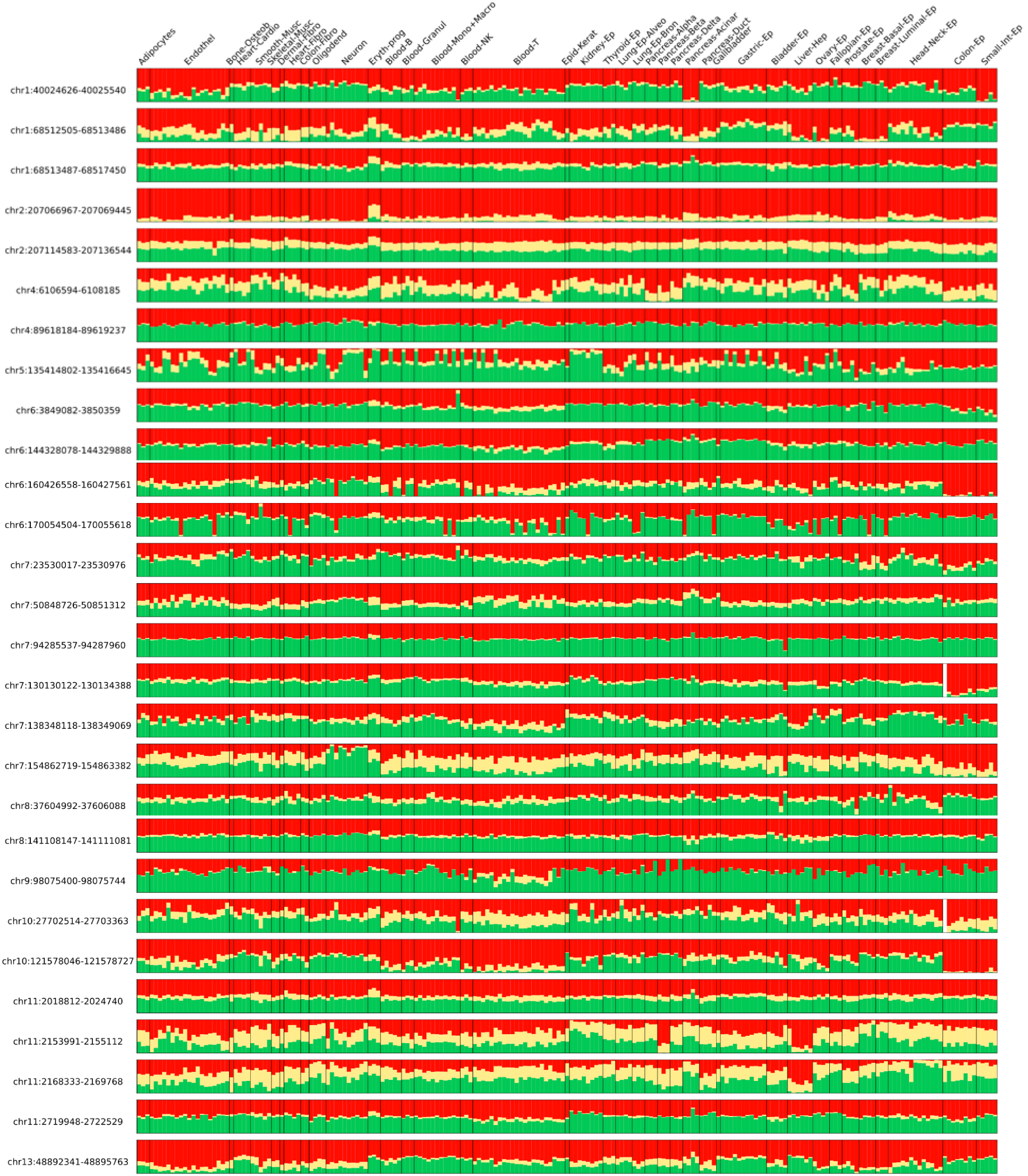
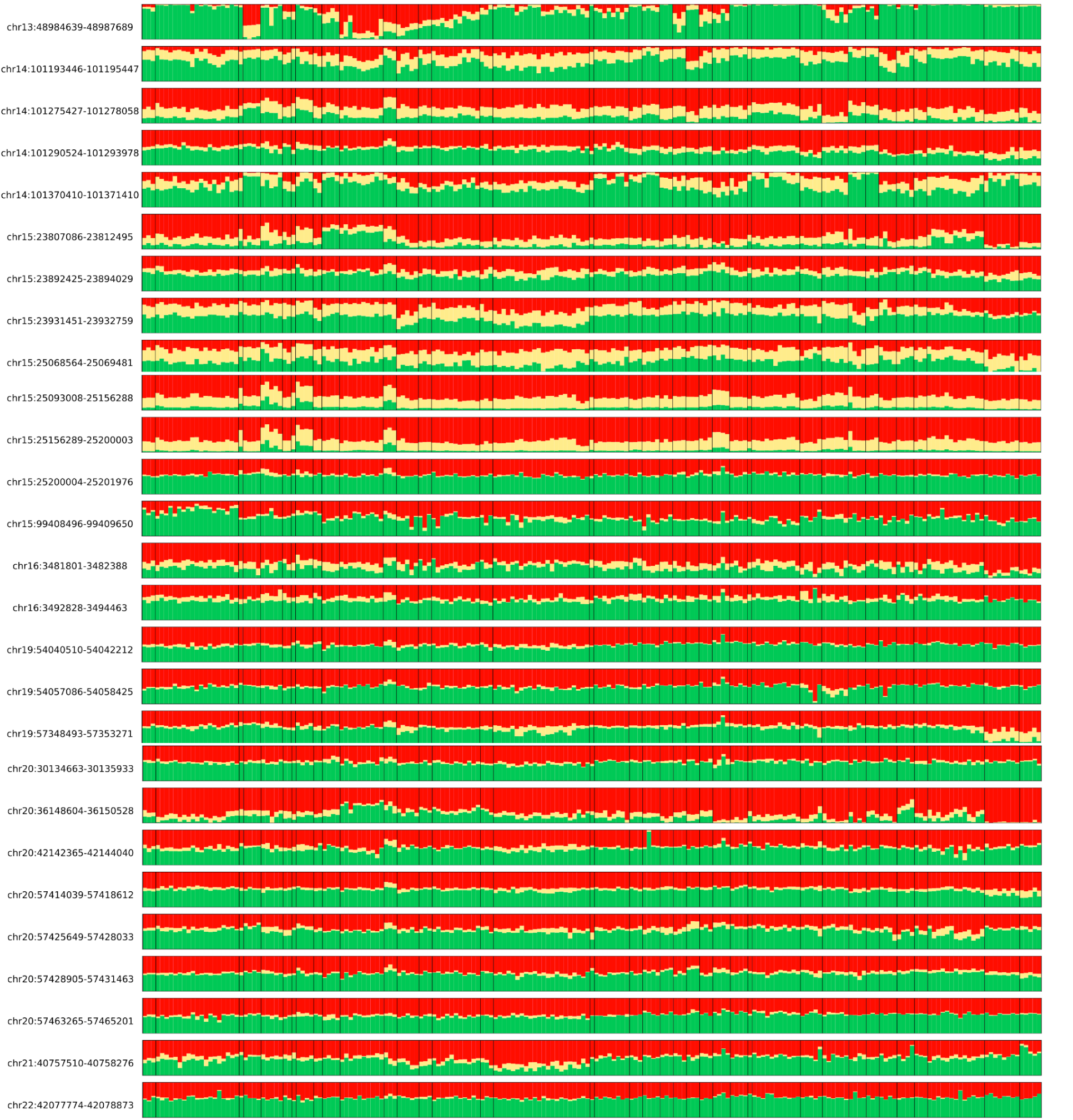
Bimodality plot of known ICRs. For each known ICR (hg19 coordinates), we plot the percent of U/X/M sequenced fragments (≥3 CpGs) across 202 samples (columns). Fully bimodal patterns are visualized is 50% unmethylated fragments (green) and 50% methylated fragments (red). Sequenced fragments with >35% unmethylated CpGs and >35% methylated CpGs (e.g. a fragment of 4 CpGs, with exactly two methylated CpGs) are classified as mixed, and their empirical percentage in a give sample/region is shown in yellow.

**Supplemental Figure 3.**
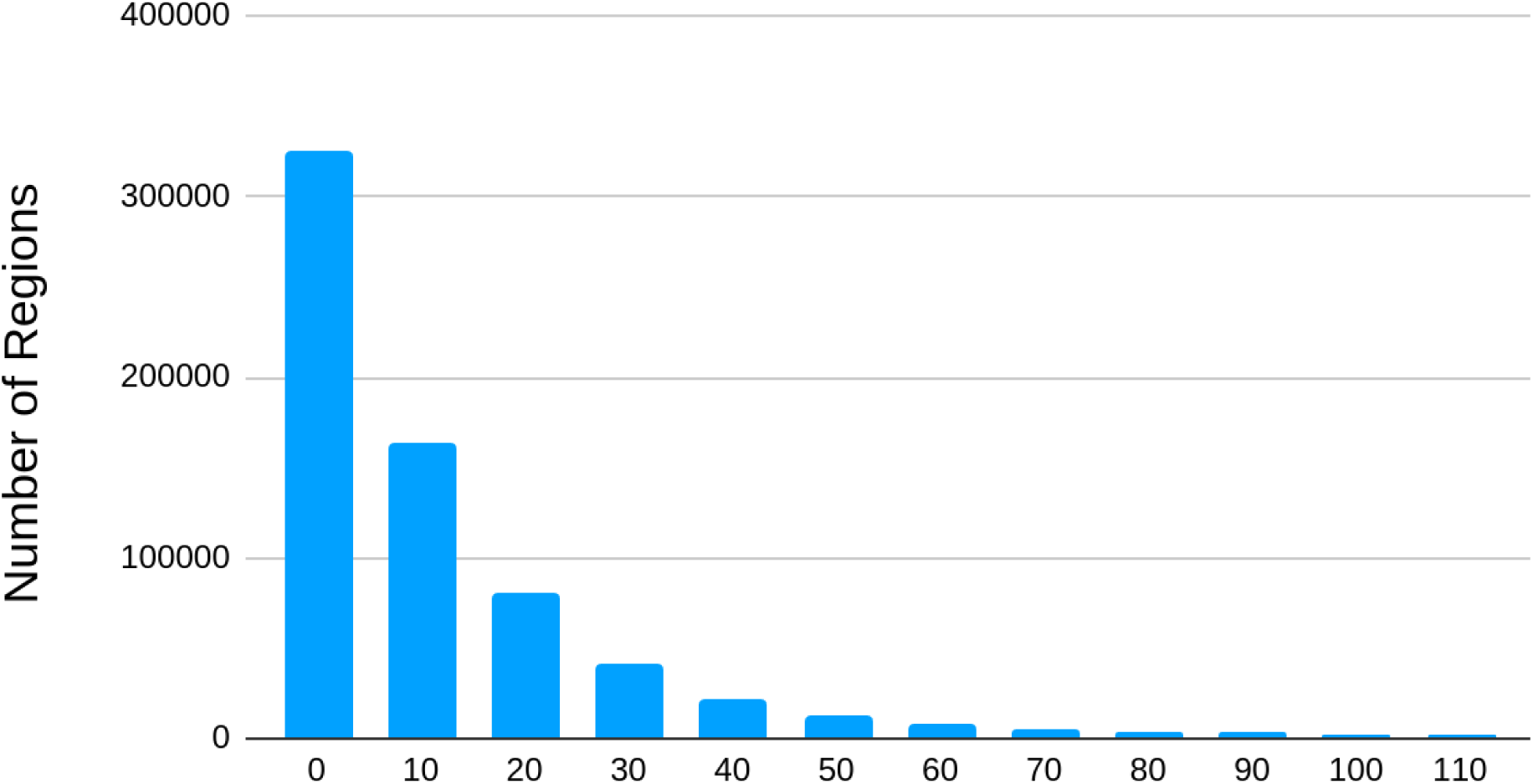
Number of regions vs number of samples exhibiting bimodal methylation. Bar chart showing the number of regions (y-axis) which are bimodal in at least the number of samples specified in the x-axis.

**Supplemental Figure 4.**
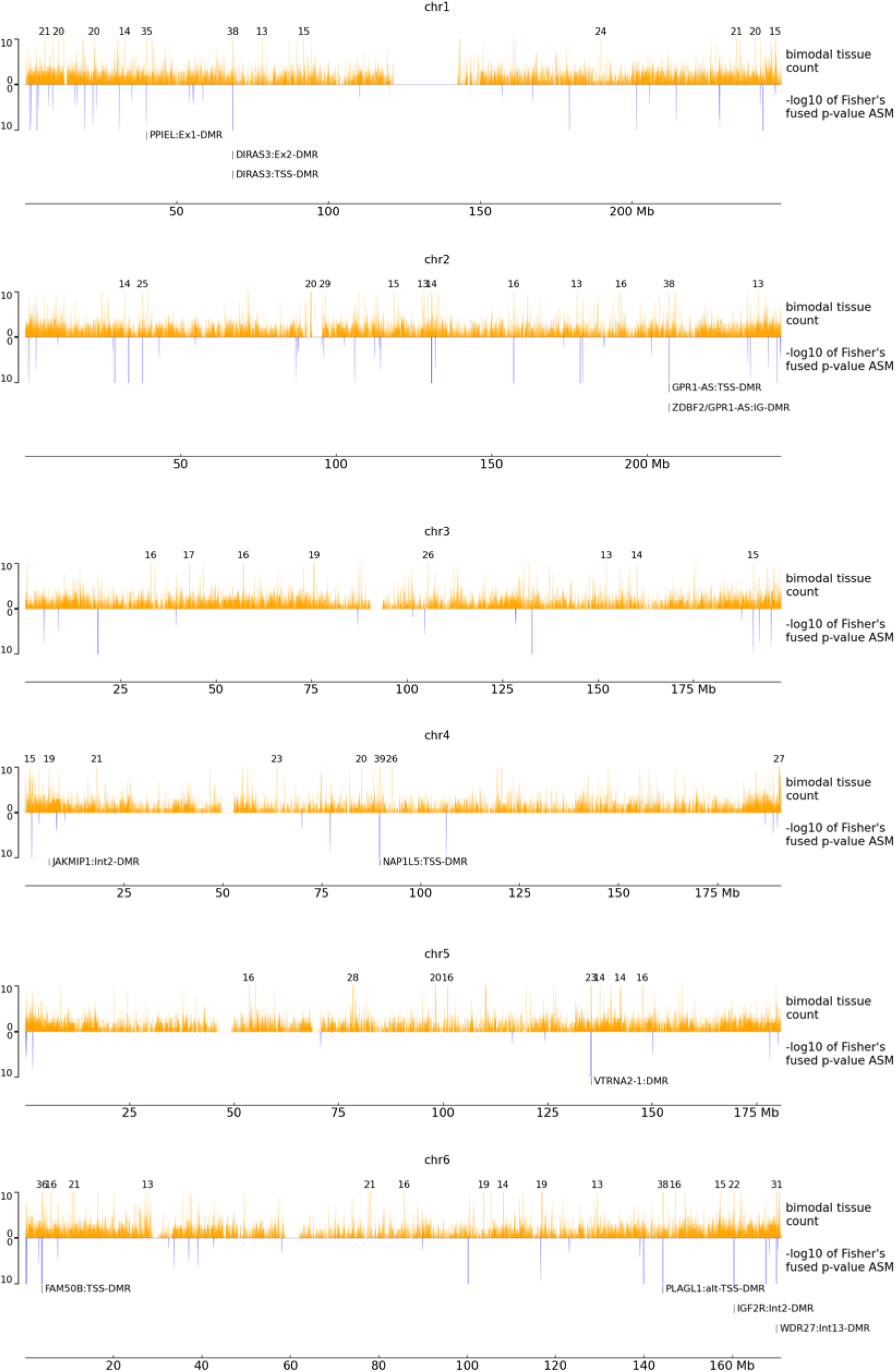
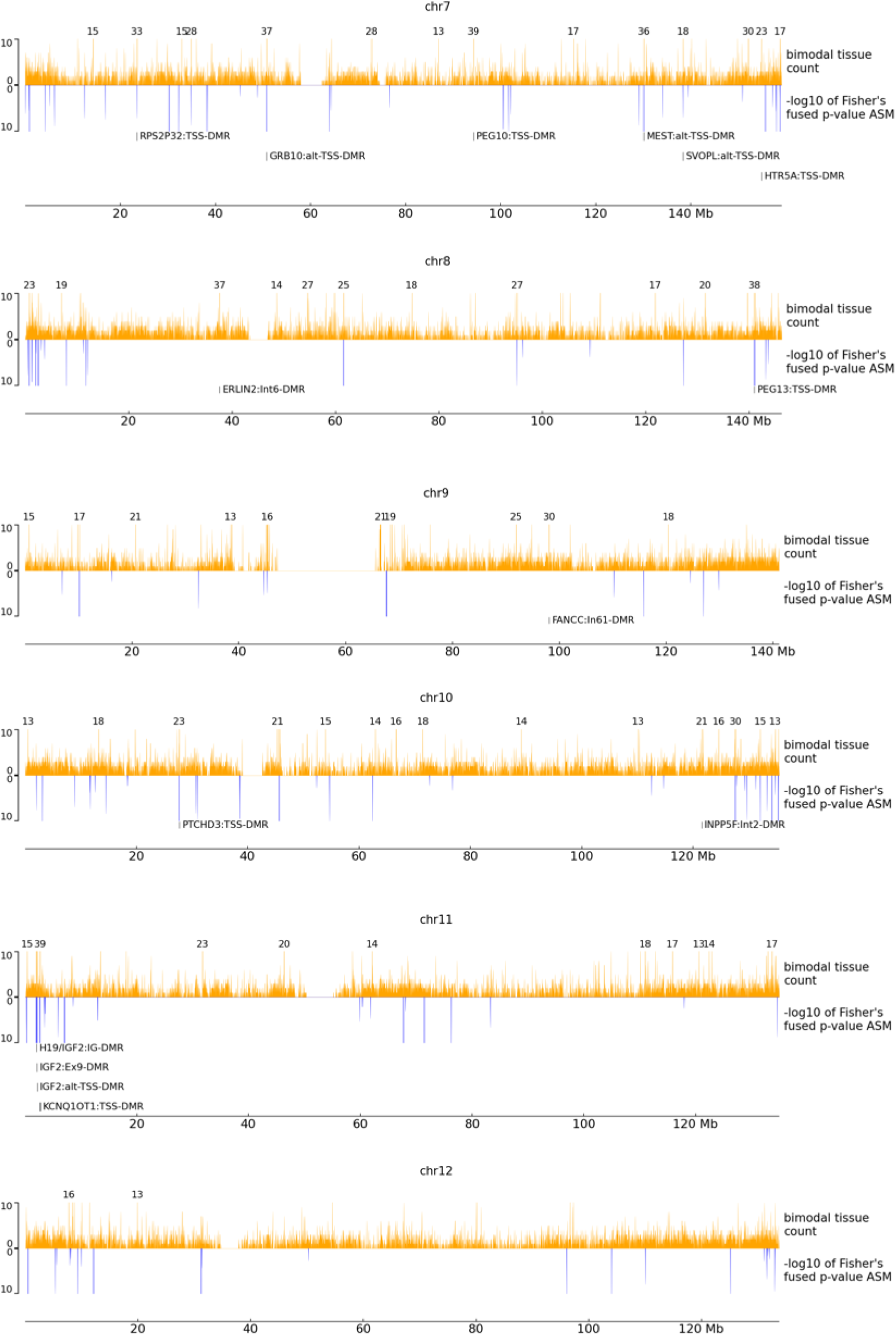
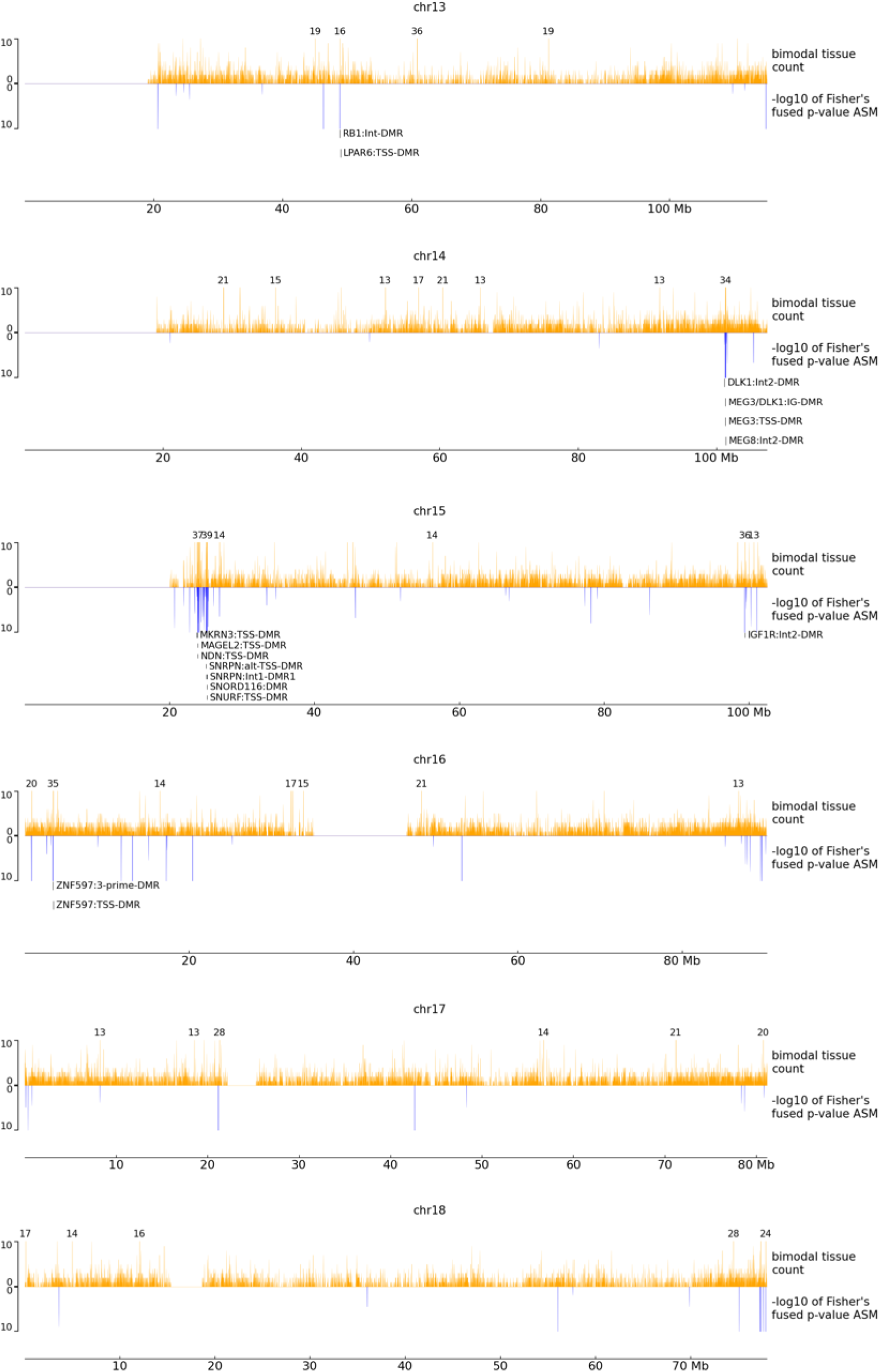
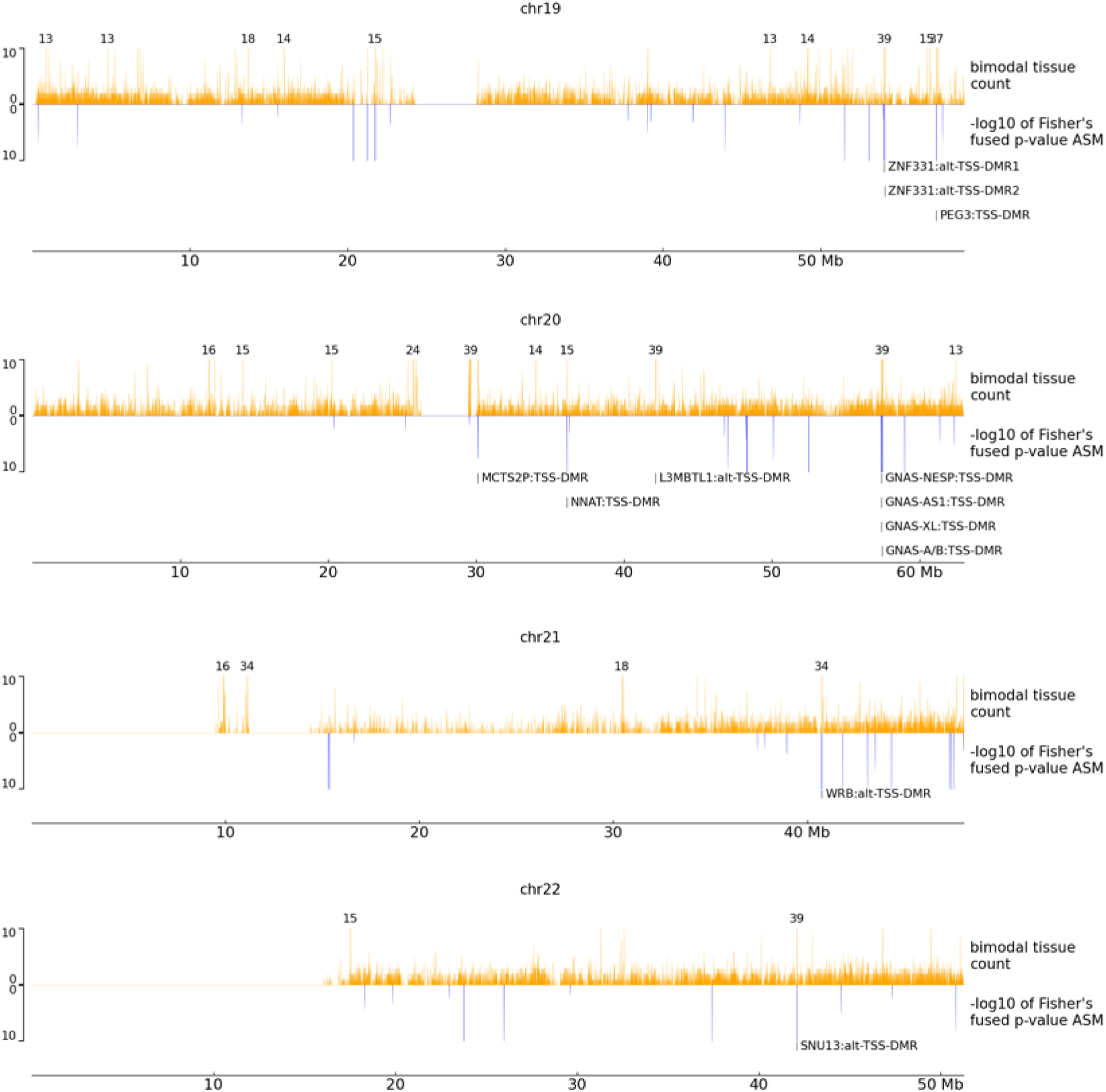
Distribution of bimodal regions and ASM across the genome. In orange, a track showing the number of tissues that are bimodal across the genome. The heights exceeding the threshold are shown above the bars. In blue the negative log10 Fisher’s fused p-value of ASM across the genome is shown. Known ICRs are indicated below each chromosome.

**Supplemental Figure 5.**
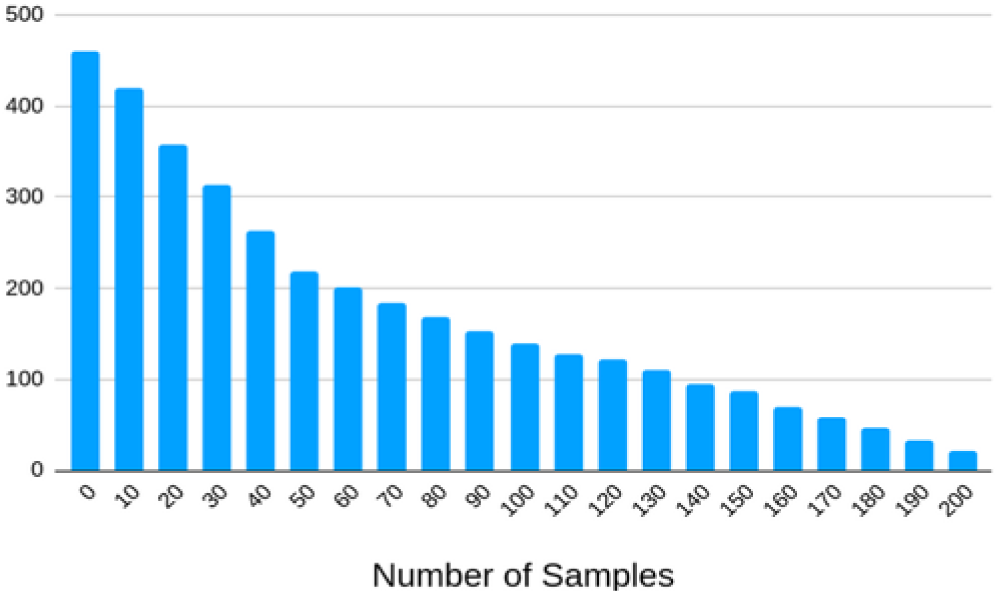
Distributional properties of parental ASM regions. Bar chart showing the number of parental-ASM-regions identified (y-axis) which are bimodal in at least the number of samples specified in the number of samples (x-axis).

**Supplemental Figure 6.**
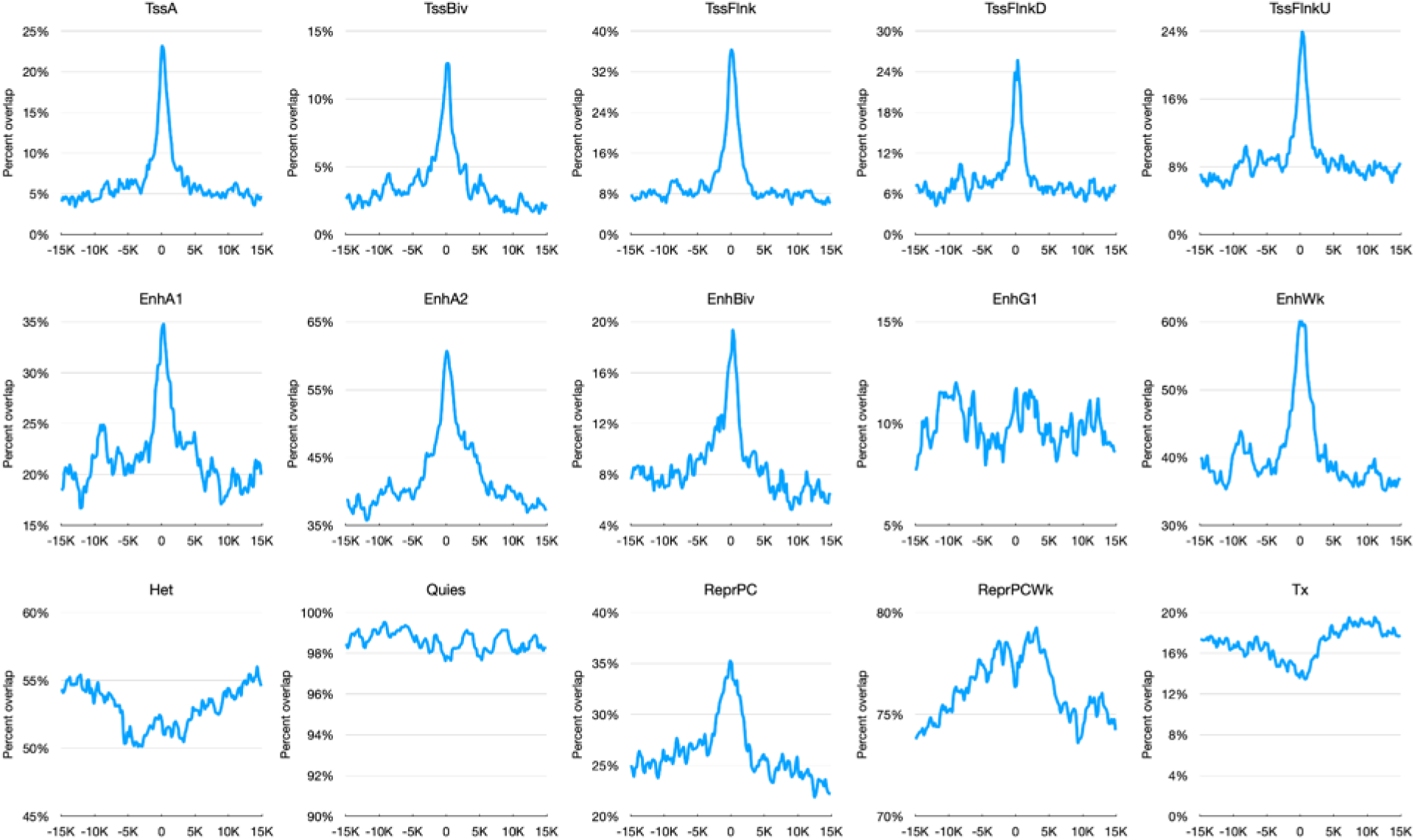
Local enrichment of parentally methylated regions for various chromHMM annotations. Average percent of intersecting nucleotides between parental-ASM regions and various chromHMM state annotations.

**Supplemental Figure 7.**
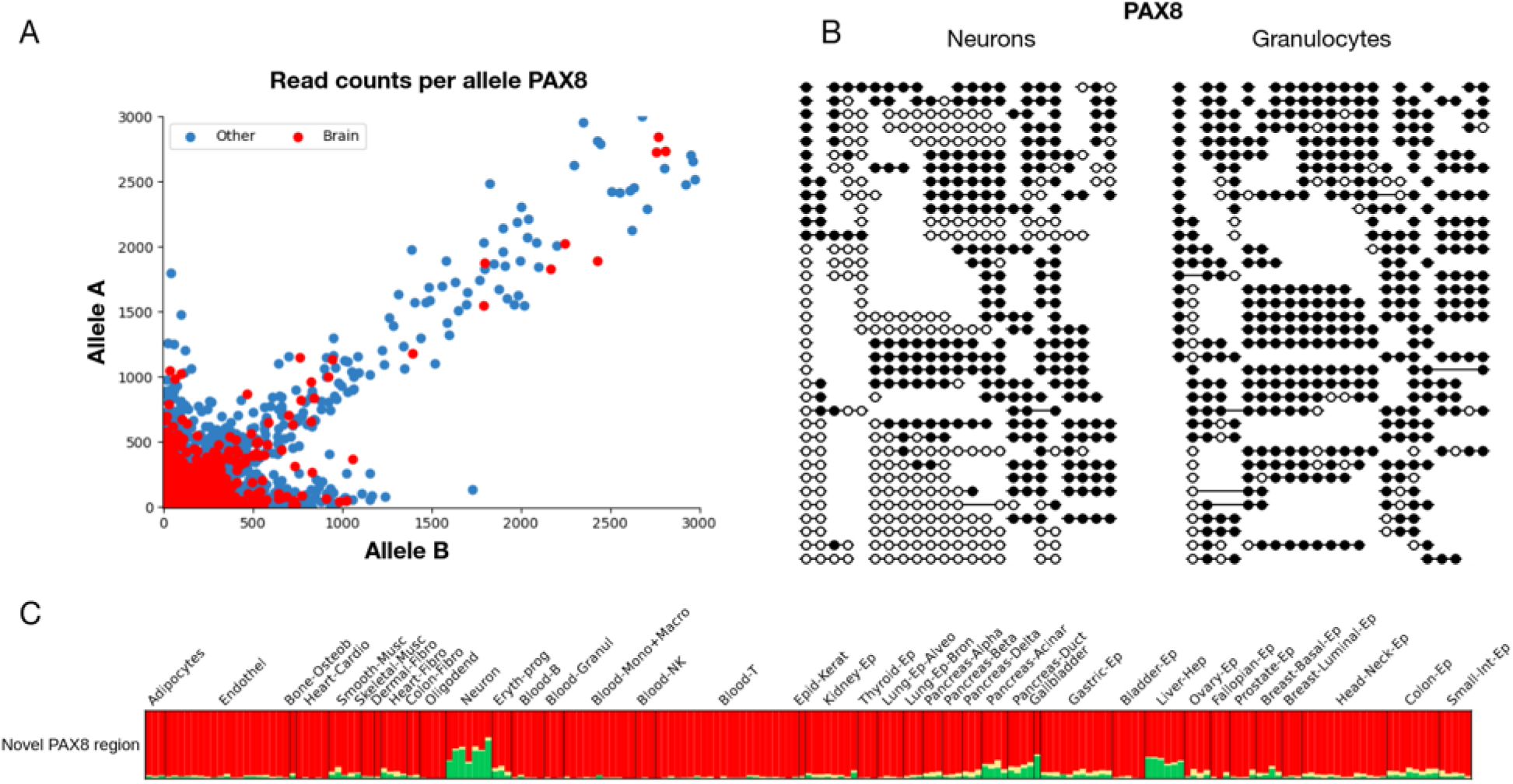
Tissue-specific allelic expression of known imprinted gene PAX8 is accompanied by matched bimodal methylation. **(A)** Allele-specific read counts from GTEx of the known imprinted gene PAX8 show imprinted expression for brain (red) samples, with other samples shown in blue. Most samples do not show allelically biased expression. Lung samples also show biased expression. **(B)** Bimodal DNA methylation in purified cortical neurons (left) but not in other samples (e.g. granulocytes, right) at a bimodal region (chr2:113953706-113955952), 17Kb upstream of PAX8). **(C)** Fragment-level methylation analysis exhibits brain-specific bimodality.

## Notes

### Competing Interest Statement

The authors have declared no competing interest.

